# Multi-Trajectory Pseudotime Inference via Permutation Factorizations

**DOI:** 10.64898/2026.05.27.728207

**Authors:** Rebecca Farina, Yikun Wang, Mariano I. Gabitto, Anamika Agrawal, Gonzalo E. Mena

## Abstract

Inferring disease progression from cross-sectional data typically relies on placing individuals along a single latent pseudotemporal axis. However, diseases often exhibit different biomarkers that evolve with different dynamics, making a single shared trajectory insufficient and often leading to biased or inconsistent orderings. We propose a probabilistic framework that maintains a shared global ordering of individuals while allowing clusters of features to follow distinct monotone trajectories along this ordering. Rather than assigning individuals to separate progressions, our approach captures heterogeneity through cluster-specific temporal responses defined with respect to a common latent sequence of individuals. Our key observation is that monotone transformations of a common progression preserve the ordering of individuals, so multiple pseudotimes factorize into a single latent permutation over individuals and cluster-specific monotonic functions, enabling flexible yet comparable representations of biomarker dynamics. We jointly infer donor ordering, biomarker clusters, and trajectories within a unified Bayesian framework, using stochastic variational inference over permutations. Across two major neuropathological datasets in Alzheimer’s disease, our method recovers biologically meaningful feature clusters and improves overall alignment with established staging measures compared to single-trajectory baselines. These results show that modeling heterogeneous dynamics relative to a shared progression yields more accurate and interpretable reconstructions of disease progression from cross-sectional data.

## 1 Introduction

Neurodegenerative diseases such as Alzheimer’s disease (AD) unfold over decades. Thus, in current paradigms, the progression of such diseases is studied from cross-sectional postmortem cohorts, which provide rich measurements of pathology, cellular states, and molecular profiles. These datasets lack an explicit temporal structure: each donor is observed only once, and reconstructing progression requires placing individuals along a latent disease trajectory before any downstream analysis can be meaningfully performed. This problem of inferring a coherent progression from heterogeneous cross-sectional observations is a central challenge in studying AD and other neurological diseases.

In current neuropathologic practice, disease burden is typically summarized using ordinal staging systems such as Braak stages [Braak and Braak, 1991] and Thal phases [Thal et al., 2002], which remain central to clinical and research assessment. However, these systems are inherently coarse: they discretize progression of AD across the brain into a small number of stages and reflect distinct, not always aligned, axes of pathology, with amyloid deposition (Thal) often preceding and partially decoupling from tau pathology (Braak) in certain regions of the brain. At the same time, recent advances in digital neuropathology and multimodal molecular profiling have made increasingly rich quantitative measurements available, creating an opportunity for data-driven models to infer continuous latent disease progression from heterogeneous cross-sectional observations spanning multiple pathological processes [Gabitto et al., 2024, Rosado et al., 2025].

A growing literature addresses this challenge by explicitly modeling a latent variable of disease progression [Campbell and Yau, 2018, 2019, Gabitto et al., 2024, Agrawal et al., 2026, Wijeratne and Alexander, 2024]. Notably, B-BIND [Agrawal et al., 2026] introduces a Bayesian framework for neurodegenerative dynamics that assigns each donor a latent pseudotime and models pathology measurements as smooth functions of this latent disease axis, thereby moving from discrete stage labels [Young et al., 2018] toward continuous progression inference in cross-sectional neuropathology. Crucially, this formulation not only yields an ordering of individuals but also enables estimation of marker-specific temporal trajectories that describe how each pathological feature evolves along the inferred disease axis, e.g., whether specific markers plateau at late stages.

Despite this progress, an important structural limitation remains: existing approaches assume that all biomarkers are functions of a single shared latent coordinate, such that each donor is assigned a single pseudotime value used uniformly across all features. While this induces a coherent global ordering, it constrains the range of temporal dynamics that can be expressed, as differences between biomarkers are limited to simple transformations of a common progression axis. However, different pathological markers may reflect a common coarse disease ordering across donors while still evolving according to partially shared, marker-specific dynamics. Amyloid- and tau-related measurements may both track disease severity, but their temporal profiles are not synchronized, and some features may be informative only over restricted portions of progression. Such heterogeneity cannot, in general, be reconciled by a single shared pseudotime, as in B-BIND [Agrawal et al., 2026]. Per-feature normalization does not resolve this: any monotone rescaling leaves each feature’s ranking of donors unchanged, and does not change the shape of its response along the latent axis, so a single parametric response cannot fit features whose dynamics differ by a nonlinear time warp. Consequently, enforcing a shared pseudotime requires the model to reconcile incompatible temporal signals across features, effectively inducing a compromise that may be dominated by a subset of biomarkers with stronger or more monotonic trends. This can lead to inferred orderings that preferentially reflect specific pathological processes while underrepresenting others, thereby failing to fully leverage the available multimodal information (see Fig. 2 for an example).

To address this limitation, we introduce **HiP-BIND** (**Hi**erarchical **P**ermutation **B**ayesian **I**nference for **N**eurodegenerative **D**ynamics), a multi-temporal hierarchical Bayesian framework for inferring disease progression from cross-sectional neuropathology data. The key idea is that, under monotonic progression, multi-feature temporal variation can be factorized into a shared global ordering of donors together with feature-specific perturbations of this ordering. This reduces the problem of inferring multiple pseudotimes to a structured permutation inference problem, enabling flexible modeling of heterogeneous trajectories with a donor ordering shared across clusters. In detail, our contributions are the following:

- **Hierarchical multi-temporal progression model**. We propose a generative model that jointly infers a global donor ordering and cluster-specific pseudotime trajectories, allowing features with heterogeneous temporal dynamics to share partial progression structure while accommodating structured deviations between clusters. Feature clustering operates in trajectory-shape space, capturing differences in growth dynamics and onset timing.
- **Relaxed permutation inference within a Bayesian framework**. We formulate donor ordering as a relaxed permutation problem embedded in a Bayesian generative model, enabling tractable posterior inference over latent orderings in a high-dimensional discrete space. We derive and implement a stochastic variational inference (SVI) scheme for scalable deployment.
- **Recovery of latent cluster structure**. HiP-BIND incorporates a clustering structure over features to capture groups of biomarkers with shared dynamics, allowing inference of interpretable feature groups directly from temporal behavior, rather than requiring groups to be specified in advance.
- **Validation on multi-region neuropathology cohorts**. We demonstrate the efficacy of HiP-BIND for analyzing two major neuropathological cohort datasets: the 84-donor SEA-AD MTG single-region dataset [Hawrylycz et al., 2024], and the ROSMAP cohort spanning eight brain regions and 1218 donors [Bennett et al., 2018]. We show that inferred cluster structure and trajectories recapitulate known disease progression patterns, including region-specific behavior and plateauing vs. sharply increasing progression for amyloid and tau markers.

HiP-BIND differs from B-BIND in three ways: (i) features are partitioned into *R* groups, each with its own monotone trajectory; (ii) the pseudotimes are factorized into a single shared donor ordering and group-specific increasing sequences coupled through a hierarchical prior; and (iii) the feature-to-group assignment is inferred jointly. With *R* = 1 the model has a single trajectory and matches B-BIND closely (Appendix B.3), so the gains we report come mainly from allowing multiple trajectory groups along the shared ordering. Equivalently, since all groups share the ordering, HiP-BIND can be read as a single donor ordering with nonparametric monotone response functions shared within each feature group; grouping pools these warps across features, which reduces the number of warps to estimate and yields the feature clusters.

## 2 Methods

Let *Y* ∈ ℕ^*M* ×*L*×*D*^ be an array of integer counts containing *M* distinct neuropathology measurements (e.g., amyloid, tau measurements) for *D* donors over *L* representations (e.g., multiple brain regions or cortical layers). We aim to simultaneously recover several targets from this matrix: 1) latent ordering of donors along a pseudotime trajectory, 2) clustering of the *M* features into *R* distinct groups with a common structure indicative of a shared biological process, and 3) feature-specific *monotonic* curves describing how each feature changes over pseudoprogression time.

To perform such inference, we consider a generative model (Fig. 1) where a first *master* pseudo-progression vector 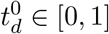, *d* ∈ [*D*], indicating donor-specific progression time, is used to define *R* distinct cluster-specific warped versions, 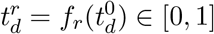, *d* ∈ [*D*], *r* ∈ [*R*], where *f*_*r*_ represents a monotonic transformation. By introducing these warped versions, we can accommodate heterogeneity in how progression may unfold across different feature groups. If we denote by *z*_*m*_ ∈ Δ^*R*−1^ the (soft) assignment of feature *m* to the *R* clusters, where Δ^*R*−1^ is the probability simplex, and write 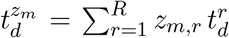 for the corresponding assignment-weighted time (equal to 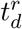 when *z*_*m*_ is the one-hot vector of cluster *r*), we can link the above specification to observations using a Negative Binomial (NB) likelihood. In detail, observations *Y*_*m,l,d*_ are independent conditional on the latents and distributed as NB with parameters 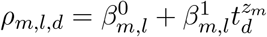 and *A*_*m,l*_:

**Figure 1:**
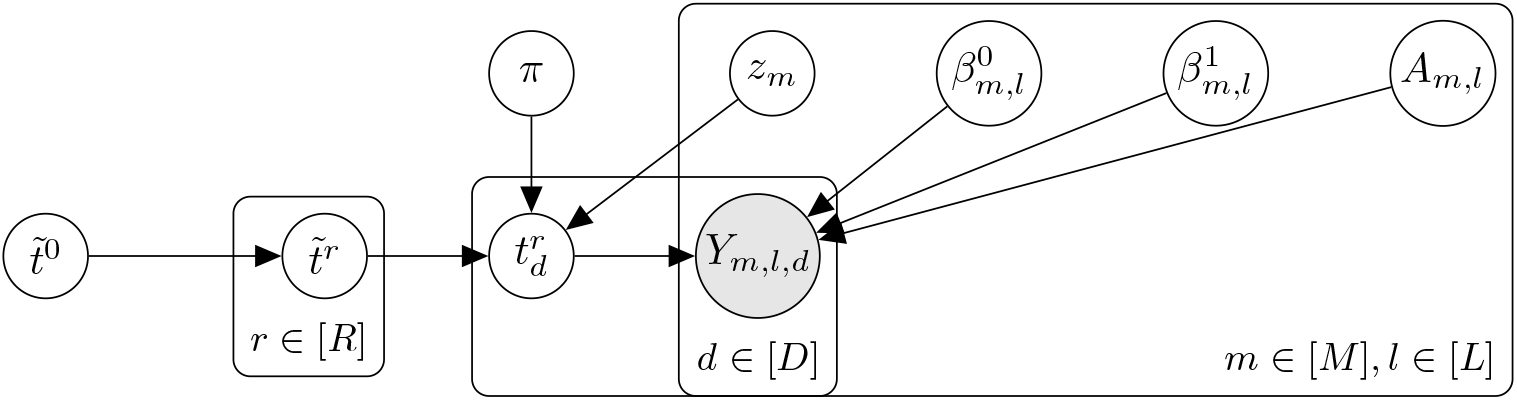
Graphical model of the HiP-BIND framework. The model factorizes progression into a global donor ordering *π* and cluster-specific monotonic trajectories, linking feature-level parameters to observed neuropathology counts.

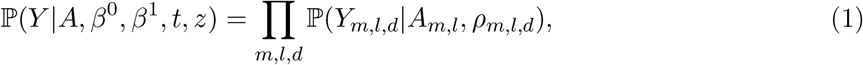

where each term is the NB probability with logits *ρ*_*m,l,d*_ and total count *A*_*m,l*_ (derived in Appendix A.1), with mean

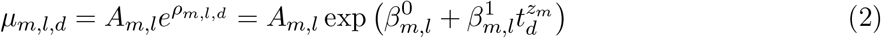

and variance 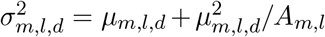. Different choices of *A, β*^0^, *β*^1^, *t* yield different curve shapes *µ*_*m,l*_ and dispersion patterns.

The model specification is completed by imposing prior distributions on the latent variables *A, β*^0^, *β*^1^, *z* and *t*. For *A, β*^0^, *β*^1^, we use hierarchical priors in the spirit of Agrawal et al. [2026] (see details in Appendix A.2). For the assignments, we use a relaxed one-hot representation, amenable to automatic differentiation: a tempered softmax of Gaussian logits, *z*_*m*_ = softmax(*u*_*m*_*/α*_*a*_) with *u*_*m*_ ~ *N* (0, *I*_*R*_) i.i.d. over *m* ∈ [*M*], which approaches a one-hot vector as the temperature *α*_*a*_ *→* 0. Our main methodological contribution is the modeling and inference of *t*, which we address in the next subsection, along with its prior specification.

### 2.1 Multiple pseudotimes with factorized permutations

The monotonicity constraint on each *f*_*r*_ is biologically meaningful but not standard in probabilistic models. We rely on two observations: first, any sequence of numbers *t* = (*t*_1_, *t*_*D*_) (not ordered) can be alternatively represented as an ordered sequence 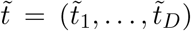, with 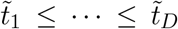, along with a *permutation*, i.e., a bijection *π* : [*D*] → [*D*] that encodes the particular ordering in which *t*_1_, …, *t*_*D*_ are presented, thus 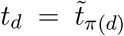. The above implies that we can specify a prior distribution over an unordered sequence *t* by specifying a prior over ordered sequences 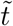 and over permutations *π*. Assuming independence among these components in the prior, we get 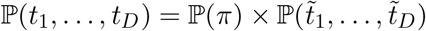. The second observation is that the constraint that all *f*_*r*_ are monotonic functions implies that whatever ordering exists in the master sequence 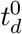, this must be preserved for the transformed 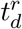, without loss of generality we take each *f*_*r*_ increasing, since decreasing responses are captured by the sign of 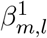. Consequently, to extend the above rationale to a joint model with multiple monotonic sequences, it suffices to encode a unique ordering *π* along with *R* + 1 increasing time sequences 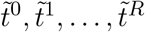, where each 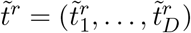 collects all donors, so that 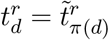. Each donor’s position along progression is encoded by *π*, shared by all clusters, while 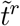 sets the spacing of progression for cluster *r*. Therefore, in this case, we can consider the following prior over unordered time sequences *t*^0^, …, *t*^*R*^ that directly extends this factorization:

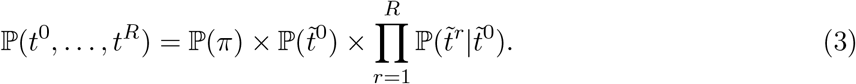

Our prior (3) is a single joint prior over all donors, not a per-donor prior, and relies on a prior specification for 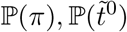 and 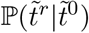. Here, we set P(*π*) ∝ 1, the uniform distribution over permutations. We impose a hierarchical structure on 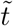: we first draw 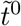 and then the remaining 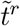 are drawn independently, conditional on this master clock. To specify priors on the sequences 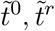 we rely on the observation that each increasing sequence with fixed endpoints, 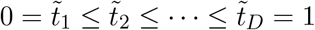, can be recovered as a cumulative sum of a zero-appended simplex-valued variable: if 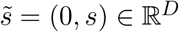 with *s* ∈ Δ^*D*−2^, then we can define 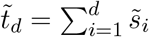. Therefore, we can define

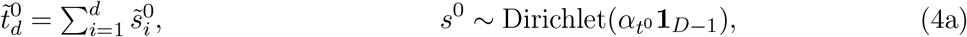

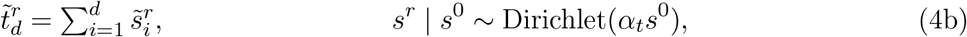

with 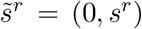 for *r* = 0, …, *R*. Note that the above prior encodes a hierarchical sampling specification: we first use the variable *s*^0^ to sample a master sequence 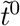 from a symmetric Dirichlet, whose concentration 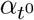 controls how far it departs from the uniform grid. Then, subsequent times 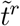 are sampled conditional on this first draw 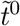, as now the Dirichlet is centered at *s*^0^, i.e., E[*s*^*r*^ *s*^0^] = *s*^0^, with *α*_*t*_ controlling the deviation. The resulting monotonic functions *f*_*r*_ are then implicit in this construction: *f*_*r*_ maps 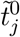 to 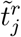 for each rank *j*. We illustrate some examples of this hierarchical specification of times in Appendix A.3.

### 2.2 Inference

Given the above specification on the likelihood and latents 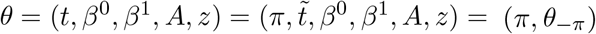 we aim to draw samples from the posterior *p*(*θ*|*Y*): computation of such posterior is intractable since it involves marginalization over *π*, which in turn involves a summation over *D*! configurations [Jerrum and Sinclair, 1989, Miller and Harrison, 2013]. To address this, we build on Linderman et al. [2018], Mena et al. [2018] and perform variational inference with two variants of the same method. In either case, we will represent *π* as a matrix *P* ∈ *B*(*D*) ⊆ ℝ^*D*×*D*^in the Birkhoff polytope (the set of doubly stochastic matrices), whose vertices are the permutation matrices (*P*_*i,j*_ = 1 iff *π*(*i*) = *j*); HiP-BIND-0 works with vertices and HiP-BIND-1 with relaxed interior points.

#### Alternating scheme (HiP-BIND-0)

We alternate between i) fixing *π* and inferring the conditional of the remaining parameters, P(*θ*_−*π*_ |*Y, π*) and ii) fixing the remaining parameters and inferring *π*, P(*π Y, θ*_−*π*_). The former is standard SVI. To sample *P*, we use the Gumbel-Matching distribution introduced in Mena et al. [2018]: given a distribution over permutations that can be written in exponential form P(*P*) ∝ exp(⟨*L, P*⟩_*F*_), we produce approximate samples by adding Gumbel noise *ε* to each entry of *L* and then produce a sample as the solution to the linear program (using the Hungarian algorithm) max_*P* ∈*B*(*D*)_ *f*⟨*P, L* + *ε*⟩_*F*_. The solution of this problem can be shown to correspond to a permutation matrix [Birkhoff, 1946]. In our case, since the likelihood factorizes over donors given 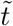, *L*_*d,j*_ is the log-likelihood of donor *d* placed at rank *j* under the current *θ*_−*π*_, so the conditional of *π* has exactly this form. We use a small Gumbel scale, so each update is effectively a maximum a posteriori step for *π*.

#### Unrolled relaxed permutation sampling (HiP-BIND-1)

We perform full variational inference over the entire parameter *θ* in a single pass. To achieve a differentiable formulation, we employ the Gumbel-Sinkhorn distribution [Mena et al., 2018] to set a prior on *B*(*D*). Iterated Sinkhorn normalization of a Gumbel-perturbed matrix makes *P* a differentiable function of noise, which enables reparameterized variational inference (details in Appendix B.1). Posterior draws of *P* are shown in Appendix Fig. 6.

We implemented all models in NumPyro [Phan et al., 2019] as hierarchical probabilistic programs that directly encode the generative model above (hyperparameters and inference settings in Appendix B.1; complexity and runtime in Appendix C). We performed posterior inference using built-in variational inference routines. This approach enables rapid model development, prototyping, and iterative refinement [Blei, 2014]. Staging labels are never used for fitting or model selection: runs and hyperparameters are selected with a label-free self-consistency criterion, the mean absolute Spearman correlation between each observed feature and the inferred ordering (Appendix B.2), which we also use to assess the choice of *R* (Appendix B.3).

As outputs, we can recover all of our targets: in addition to *t*^0^ representing the master progression timing and *z* representing feature clustering, we recover the mean vectors *µ*_*m,l*_ ∈ ℝ^*D*^ in Eq. (2). By plotting these curves as functions of *t*^0^, we can represent how each pathology marker evolves over time. Finally, we note the inherent lack of identifiability in our formulation: although the shared permutation structure ensures that the relative ordering of donors along *t*^0^ is identifiable, the absolute geometry of this coordinate is not. Monotonic transformations of *t*^0^ can be offset by corresponding changes in the cluster-specific warping functions and feature-level parameters, yielding equivalent likelihoods. Consequently, *t*^0^ is best interpreted as a canonical latent coordinate that aligns heterogeneous feature trajectories, rather than a direct estimate of absolute progression time or population sampling density.

## 3 Simulations

To isolate the effect of heterogeneous temporal dynamics, we construct a synthetic setting where features share the same mean function but differ only through temporal warping. Specifically, we consider a negative binomial setting with 100 donors and two latent monotone trajectories, where features 1–5 follow a logistic time warp, while features 6–10 follow a power-law warp. The two groups share the same mean-function coefficients but differ by a nonlinear time warp, making temporal alignment the primary challenge. Additional simulations with alternative warping functions and with uninformative features are provided in Appendix E. We fit HiP-BIND with two latent clusters, matching the true number of feature groups. To mimic the real cross-sectional setting, all methods are fit to the unordered observed count data, without access to the true donor ordering, which is used only for evaluation. We run each model across multiple seeds, which affect initialization and stochastic optimization, and report results over 20 independent simulation replicates to assess robustness rather than relying on a single run. HiP-BIND is implemented using the variational inference procedure described in Section 2.2, and performance is evaluated by Spearman correlation between inferred and true latent times, which measures agreement in donor ranking, and by Mallows distance between inferred and true permutations [Mallows, 1957]. We also fitted HiP-BIND with *R* = 1, …, 5. Under-specifying *R* (*R* = 1) sharply degrades recovery, whereas all *R* ≥ 2 perform well, and a label-free self-consistency criterion recovers the true *R* = 2 in all 20 datasets (see Appendix B.3 for details).

Fig. 2 (top) shows the inferred mean trajectories plotted against the recovered donor ordering. Both HiP-BIND models recovered the distinct latent warping functions. In contrast, the B-BIND baseline fails to capture the second trajectory. The successful trajectory recovery by HiP-BIND is driven by highly accurate latent donor ordering; the inferred permutation matrices for the order-aware models exhibit a near-diagonal structure closely matching the ground truth (Fig. 2, bottom), whereas the baselines exhibit substantial off-diagonal mass. Quantitatively, the HiP-BIND models achieved higher Spearman correlations and lower Mallows distances (Table 1; paired tests in Appendix D.1) than the B-BIND models, confirming the accurate alignment and trajectory recovery visually evident in Fig. 2. These results suggest that explicitly modeling ordering and assignment is critical when heterogeneity arises through temporal warping rather than through changes in the mean function itself, and this cannot be captured by a single shared pseudotime. PCA outperforms B-BIND in this setting because B-BIND must explain both time warps with a single monotone response, so its ordering is pulled toward one feature group, whereas the leading principal component of the standardized features combines both groups and tracks their shared axis. While PCA achieves moderately high Spearman correlations, the resulting PCA trajectories in Fig. 2 do not preserve the two monotone temporal processes in a biologically interpretable way. PCA is an unconstrained linear projection and therefore does not enforce the monotone progression structure required by our modeling goal. Since our focus is on models that infer coherent monotone disease progression, we exclude PCA from further analyses.

**Figure 2:**
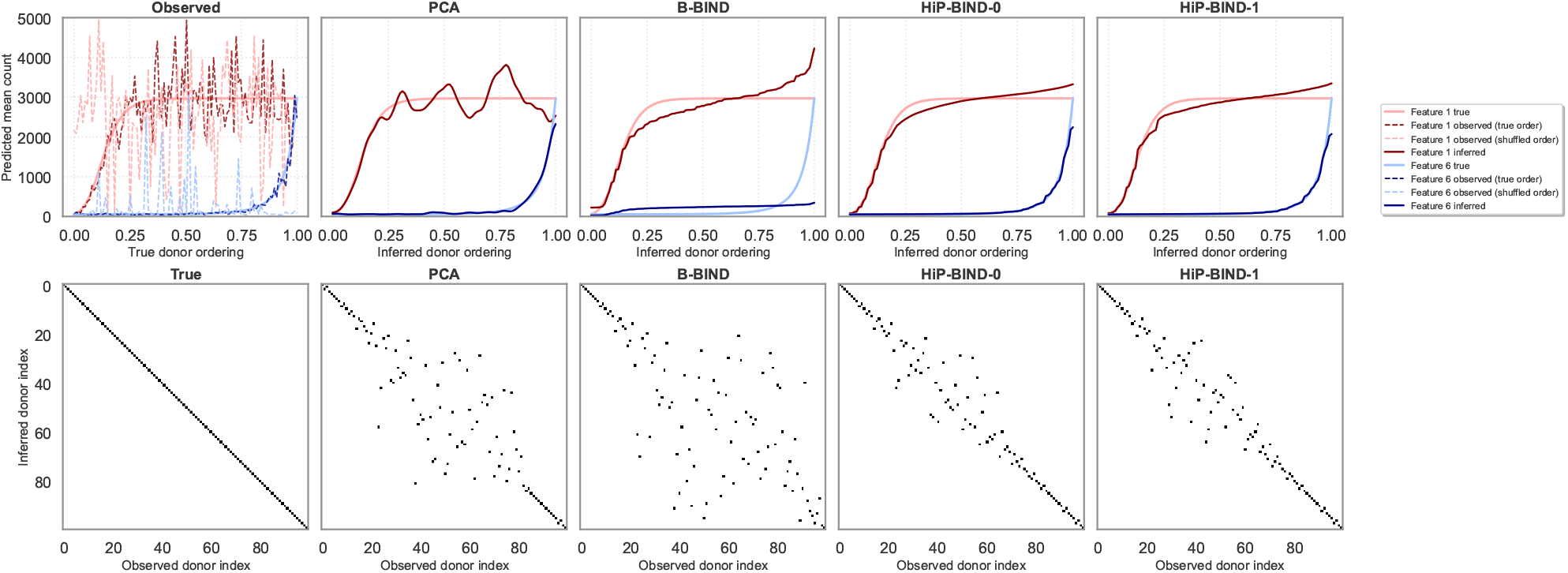
(Top) Predicted mean feature count trajectories plotted against the inferred donor ordering. Each curve corresponds to a feature, with colors indicating features generated from two distinct latent temporal processes: red curves (feature 1) follow a logistic time warp, while blue curves (feature 6) follow a power-law warp. Solid lines denote the inferred trajectories. In the left-most panel, dashed lines show the observed data displayed both in the true donor order and in the shuffled order used for model fitting; all methods are fit only to the shuffled, unordered observations and do not have access to the true ordering. (Bottom) Inferred permutation matrices representing the recovered latent donor orderings (true vs. inferred donor index); a diagonal structure indicates accurate recovery.

**Table 1:** Spearman correlation between the inferred and true latent times and Mallows distance for the inferred permutation matrices, averaged across 20 independent runs. Higher Spearman and lower Mallows are better. Paired significance tests are in Appendix D.1.

| Method | Spearman (mean ± std) | Mallows (mean ± std) |
| --- | --- | --- |
| PCA | 0.8947 ± 0.0190 | 13.1951 ± 1.2011 |
| B-BIND | 0.7292 ± 0.0450 | 21.1695 ± 1.7990 |
| HiP-BIND-0 | 0.9669 ± 0.0076 | 7.3778 ± 0.8342 |
| HiP-BIND-1 | **0.9670** ± 0.0089 | **7.3541** ± 0.9965 |

## 4 Results on Real Data

### 4.1 Data description

We evaluate HiP-BIND on two cross-sectional postmortem neuropathology datasets spanning complementary scales of Alzheimer’s disease pathology. The first dataset is the Seattle Alzheimer’s Disease Brain Cell Atlas (SEA-AD) [Hawrylycz et al., 2024, Gabitto et al., 2024]. SEA-AD contains quantitative neuropathology measurements from the middle temporal gyrus (MTG) of 84 postmortem brain donors. The MTG is measured across cortical layers, and the available features include amyloid-beta burden measured by 6e10 immunoreactivity, hyperphosphorylated tau measured by AT8 immunoreactivity, neuronal marker NeuN, and microglia-amyloid colocalization features. We treated features as the *M* dimension in our model, and layers (SEA-AD) and regions (ROSMAP) as the secondary *L* axis. In our analysis, we use the subset of SEA-AD markers with sufficient signal and coverage across donors, following the feature-selection strategy used in Agrawal et al. [2026].

The second dataset is a multi-region neuropathology cohort derived from the Religious Orders Study and Rush Memory and Aging Project (ROSMAP) [Bennett et al., 2018], which combines longitudinal clinical characterization with postmortem neuropathological assessment and brain donation. In contrast to SEA-AD, which is restricted here to MTG, the ROSMAP neuropathology data used in our experiments contain pathology measurements across eight brain regions, including entorhinal cortex (EC), hippocampus (Hip), angular gyrus (AG), and inferior temporal cortex (IT). We include 1218 donors. We analyze quantitative measures of amyloid-beta burden, tau pathology, neurofibrillary tangles, neuritic plaques, and diffuse plaques; amyloid and tau measurements are analyzed on the square-root scale, as provided in the source data. For both SEA-AD and ROSMAP, data access, donor inclusion criteria, the mapping of measurements to counts, and a complete list of markers are given in Appendix F.

We used these two datasets to evaluate (i) how well HiP-BIND can integrate information from different features and improve donor ordering along progression, (ii) whether the recovered trajectories reflect known progression growth patterns, and (iii) whether HiP-BIND-induced feature clusters capture meaningful biological structure.

### 4.2 Integration of different markers and progression-order inference

We aim to determine if HiP-BIND is able to recover progression order in a way that integrates information from different features. Since ground-truth progression ordering is unavailable in cross-sectional postmortem data, we evaluated it by analyzing correlation patterns with standard ordinal staging measures: Braak stage, Thal phase, CERAD score [Mirra et al., 1991, Rossetti et al., 2010], and ADNC score [Hyman et al., 2012]. These clinical and neuropathological scores summarize partially overlapping but distinct axes of Alzheimer’s disease progression and, therefore, provide a useful test of whether an inferred pseudotime captures a broad disease signal rather than a single marker-specific process. Specifically, Braak measures tau pathology, Thal measures amyloid pathology, CERAD measures neuritic plaque density, and ADNC is a composite score incorporating amyloid, tau and neuritic plaque burden. Therefore, in the absence of ground truth, we view ADNC as a composite measure of success in recovering a pseudo-ordering that integrates amyloid- and tau-related processes.

We fit HiP-BIND-0 with *R* = 3 latent trajectory clusters of features to allow the model to represent interpretable trajectory groups, approximately corresponding to amyloid-dominant, tau-dominant, and mixed or region-specific temporal patterns. With *R* = 1, HiP-BIND performs like the all-marker B-BIND, whereas *R* = 2, …, 5 give similar self-consistency scores and staging correlations (Appendix B.3). We compared HiP-BIND to B-BIND fitted to the same full set of markers, as well as B-BIND variants fit only to amyloid markers, only to tau markers, or to amyloid and tau markers. These variants allow us to assess the extent to which single-trajectory models can capture different aspects of disease progression when driven by specific marker classes. We report average Spearman correlations over 20 runs per model (selection in Appendix B.2) in Table 2, with standard deviations and statistical significance tests in Appendix D.2. For a more robust assessment, we also stratify the analysis by APOE status [Liu et al., 2013].

**Table 2:** Mean absolute Spearman correlation between model-inferred donor orderings and standard ordinal clinical scores (Braak, Thal, ADNC, and CERAD), stratified by APOE carrier status (All, APOE+, APOE-). Results are averaged across 20 runs. Standard deviations and significance tests are in Appendix D.2.

| (a) ROSMAP |  |  |  |  |  |  |  |  |  |  |  |  |
| --- | --- | --- | --- | --- | --- | --- | --- | --- | --- | --- | --- | --- |
| Method | All |  |  |  | APOE+ |  |  |  | APOE- |  |  |  |
|  | Braak | Thal | ADNC | CERAD | Braak | Thal | ADNC | CERAD | Braak | Thal | ADNC | CERAD |
| HiP-BIND-0 | 0.740 | 0.842 | **0.883** | **0.836** | 0.808 | **0.677** | **0.816** | **0.652** | 0.699 | 0.861 | **0.888** | **0.863** |
| B-BIND (All markers) | 0.648 | **0.844** | 0.828 | 0.816 | 0.624 | 0.643 | 0.665 | 0.600 | 0.626 | **0.868** | 0.851 | 0.848 |
| B-BIND (Amyloid + Tau) | 0.634 | 0.821 | 0.809 | 0.780 | 0.587 | 0.601 | 0.626 | 0.540 | 0.616 | 0.847 | 0.832 | 0.812 |
| B-BIND (Amyloid) | 0.576 | 0.809 | 0.780 | 0.763 | 0.492 | 0.568 | 0.547 | 0.494 | 0.557 | 0.840 | 0.813 | 0.800 |
| B-BIND (Tau) | **0.788** | 0.590 | 0.710 | 0.596 | **0.828** | 0.590 | 0.789 | 0.565 | **0.763** | 0.564 | 0.664 | 0.575 |

| (b) SEA-AD |  |  |  |  |  |  |  |  |  |  |  |  |
| --- | --- | --- | --- | --- | --- | --- | --- | --- | --- | --- | --- | --- |
| Method | All |  |  |  | APOE+ |  |  |  | APOE- |  |  |  |
|  | Braak | Thal | ADNC | CERAD | Braak | Thal | ADNC | CERAD | Braak | Thal | ADNC | CERAD |
| HiP-BIND-0 | 0.710 | **0.843** | **0.814** | 0.665 | 0.485 | **0.624** | 0.458 | 0.234 | 0.701 | **0.839** | **0.829** | **0.737** |
| B-BIND (All markers) | 0.650 | 0.828 | 0.769 | 0.625 | 0.433 | 0.603 | 0.380 | 0.153 | 0.624 | 0.822 | 0.780 | 0.704 |
| B-BIND (6e10 + AT8) | 0.657 | 0.811 | 0.776 | 0.618 | 0.458 | 0.585 | 0.404 | 0.158 | 0.633 | 0.801 | 0.779 | 0.675 |
| B-BIND (6e10) | 0.637 | 0.824 | 0.755 | 0.623 | 0.408 | 0.593 | 0.349 | 0.124 | 0.615 | 0.828 | 0.778 | 0.719 |
| B-BIND (AT8) | **0.787** | 0.685 | 0.794 | **0.672** | **0.738** | 0.463 | **0.693** | **0.608** | **0.739** | 0.628 | 0.743 | 0.621 |

Across both ROSMAP and SEA-AD, HiP-BIND achieves the strongest overall agreement with established staging measures (Table 2). In ROSMAP, it attains the highest average correlations for ADNC and CERAD, and remains competitive for Braak and Thal, where it is outperformed only by the single-marker tau B-BIND (Braak) and, marginally, by the all-marker B-BIND (Thal). Under APOE stratification, HiP-BIND remains strongest or competitive across all staging measures, and its gains over the all-marker B-BIND are largest among APOE carriers. The same trend holds in SEA-AD: HiP-BIND outperforms multi-marker B-BIND across all staging measures in the full cohort, attaining the highest correlation of all methods for Thal and ADNC. Paired donor-bootstrap analyses showed that the improvements over all-marker B-BIND were statistically supported for Braak, ADNC, and CERAD in both cohorts, whereas the differences for Thal were not statistically distinguishable from zero (Appendix Table 8).

As expected, tau-only B-BIND aligns most strongly with Braak staging, while amyloid-only B-BIND aligns most strongly with Thal staging, reflecting the pathology-specific nature of these metrics. However, such limited baselines typically lead to poor scores in the remaining metrics. Additionally, in both datasets, correlations for B-BIND fitted to all markers closely track the ones for B-BIND fitted to amyloid only features. This result is consistent with the simulation in Fig. 2, where B-BIND effectively tracks only one of the feature groups, indicating that it does not fully integrate tau information. Thus, B-BIND can capture one dominant pathological signal at a time, but not both simultaneously. HiP-BIND overcomes this limitation by maintaining a shared ordering while allowing marker-specific temporal variation. These findings support the central premise of the model: allowing marker groups to follow distinct temporal responses improves recovery of a shared donor ordering across heterogeneous pathological features.

### 4.3 Clustering

Fig. 4 highlights an additional advantage of HiP-BIND: it recovers biologically meaningful feature structure directly from temporal dynamics. In both datasets, amyloid- and tau-related features form distinct clusters, consistent with their known differences in disease progression. In particular, in ROSMAP, amyloid, diffuse plaques and neuritic plaques form one cluster and tau and tangles another. In SEA-AD, the 6e10 and Iba1–6e10 features cluster together, NeuN joins them at a larger distance, and the two AT8 features form a separate cluster. The clustering is also stable across related measurements. This structure is not available in single-trajectory models and demonstrates that multi-trajectory inference improves not only ordering but also interpretability of the underlying pathology.

To further examine regional structure in the multi-region ROSMAP cohort, we fitted HiP-BIND to a flattened data matrix with dimensions *M*′ = *M* × *L* features and *L*′ = 1 layers, treating layers as pseudo-features. While amyloid measurements from different regions remained grouped within the same cluster, tau measurements separated into two distinct clusters (see Appendix Fig. 12). One cluster contained entorhinal cortex and hippocampus, consistent with the well-established regional heterogeneity of tau dynamics: tau pathology emerges early in the medial temporal lobe before spreading to neocortical regions at later stages of disease [Braak and Braak, 1991, Berron et al., 2021, Adams et al., 2022, Doering et al., 2024].

### 4.4 Recovered trajectories

Fig. 3 provides a trajectory-level view of this behavior, comparing HiP-BIND-0 with the all-marker B-BIND baseline. For visualization, we consider a subset of all markers, and each plot uses the run whose Spearman correlation is closest to the median across 20 seeds for the corresponding dataset–model combination. Trajectories for all features are shown in Appendix G (Figs. 10 and 11). In ROSMAP, amyloid trajectories increase smoothly across regions, while tau exhibits more region-specific and often sharper late-stage dynamics. These differences cannot be reconciled by a single temporal response, forcing B-BIND into a compromise. HiP-BIND instead captures these processes as distinct but coordinated trajectories. The same pattern appears in SEA-AD, where amyloid (6e10) increases gradually, while tau (AT8) remains flat for much of the ordering before rising sharply at later stages. This mismatch in temporal profiles is not well captured by a single pseudotime, but is naturally accommodated by HiP-BIND.

**Figure 3:**
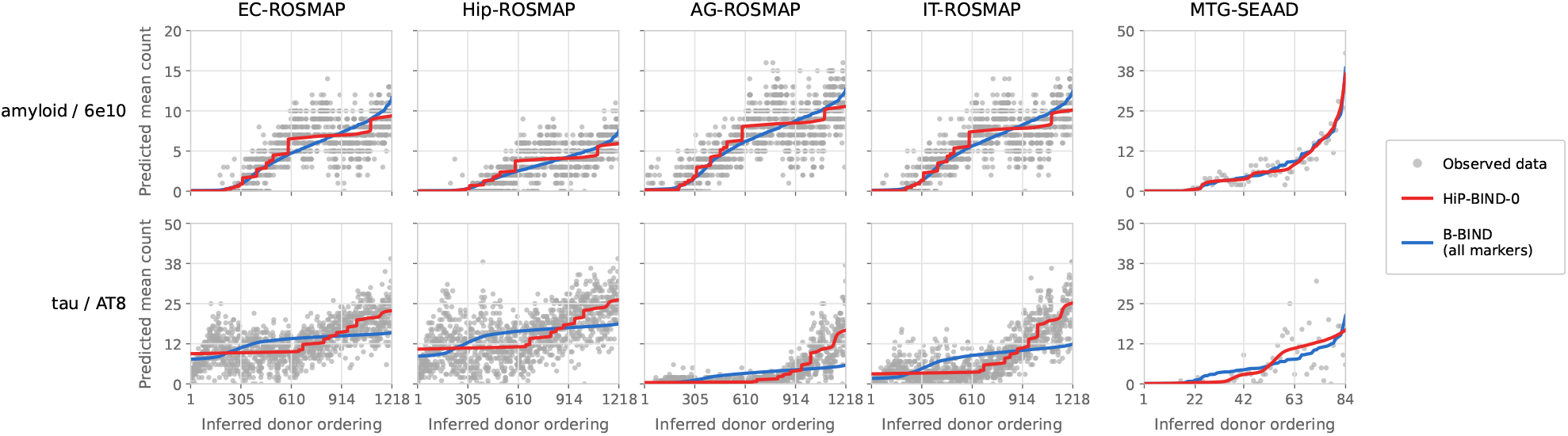
Feature-specific trajectory reconstruction. Predicted monotonic feature trajectories plotted against each model’s inferred donor ordering, with observed donor measurements overlaid using the HiP-BIND-0 inferred ordering. Columns 1–4: amyloid and tau in four ROSMAP regions (EC, Hip, AG, IT); column 5: amyloid (percent 6e10 positive area) and tau (percent AT8 positive area) in the SEA-AD MTG region.

**Figure 4:**
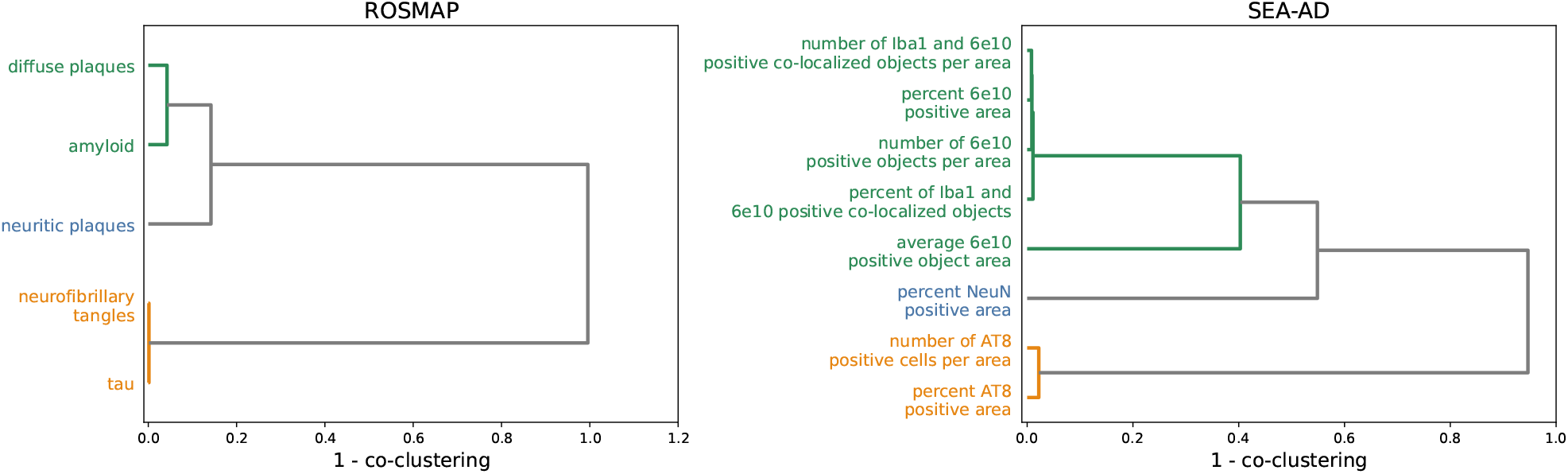
Latent feature clustering recovers known biological relationships. Dendrograms illustrating the hierarchical clustering of neuropathological features based on their inferred temporal dynamics via HiP-BIND-0 for the ROSMAP (left) and SEA-AD (right) cohorts. Distance is measured as 1 − *P* (co-cluster), where *P* (co-cluster) is the soft co-clustering probability averaged across the 20 runs. The recovered clusters separate amyloid- and tau-related features, indicating that HiP-BIND identifies biologically meaningful groups from inferred temporal dynamics.

## 5 Conclusions

We introduced HiP-BIND, a hierarchical Bayesian framework for multi-trajectory pseudotime inference from cross-sectional neuropathology data. By factorizing disease progression into a shared donor ordering and cluster-specific monotone trajectories, the model preserves a common notion of progression while allowing heterogeneous biomarker dynamics. In simulations, HiP-BIND recovered both the latent donor ordering and distinct temporal warping patterns more accurately than single-trajectory baselines. In SEA-AD and ROSMAP, the inferred orderings showed strong agreement with established neuropathological staging measures, while the recovered trajectories and feature clusters reflected known amyloid- and tau-related progression patterns. Together, these results suggest that modeling heterogeneous temporal responses relative to a shared progression axis can improve both donor ordering and biological interpretability.

Despite these promising results, our framework has some limitations. First, the model assumes that each feature’s mean is monotone along its group’s pseudotime (the noisy observations need not be monotone), which may not fully capture biomarkers exhibiting transient or non-monotone behavior; the group-specific warps are conditionally independent given the master clock *t*^0^, so dependence between groups beyond the shared ordering is not modeled. Second, the inferred pseudotime represents a relative latent ordering rather than an absolute estimate of disease time, reflecting the inherent non-identifiability of progression geometry from cross-sectional data alone. Finally, without longitudinal observations, inference cannot establish causal temporal progression. Although we evaluate HiP-BIND on Alzheimer’s disease, none of its components is AD-specific: it requires only cross-sectional, non-negative markers whose means change monotonically along a shared progression. It could therefore be applied to other progressive disorders in which several pathological processes co-evolve with distinct dynamics, such as Huntington’s disease, amyotrophic lateral sclerosis or progressive supranuclear palsy, and, with a different likelihood, to continuous in-vivo biomarkers. Extending the analyses to these diseases, richer trajectory classes and integration with longitudinal measurements are left for future work.

## Acknowledgements

A.A. acknowledges support from the Shanahan Foundation Fellowship. M.I.G. was supported by National Institutes on Aging (NIA) Grant U19AG060909 and R01102-01-071-20. G.M. was supported by NSF DMS-2412895. This work used Bridges-2 at Pittsburgh Supercomputing Center through allocation MTH230027 from the Advanced Cyberinfrastructure Coordination Ecosystem: Services & Support (ACCESS) program, which is supported by NSF Grant 2138259, 2138286, 2138307, 2137603, and 2138296. ROSMAP is supported by P30AG10161, P30AG72975, R01AG17917, R01 AG015819, U01 AG072572, and U01 AG046152. ROSMAP resources can be requested at https://www.radc.rush.edu and https://www.synapse.org. For correspondence, please contact G.M. and A.A. at the listed email addresses.

## Ethical approval

All participants enrolled in ROSMAP without known dementia and agreed to detailed clinical evaluation and brain donation at death. All studies were approved by an Institutional Review Board of Rush University Medical Center. Each participant signed informed and repository consents and all ROSMAP participants signed an Anatomic Gift Act. ROSMAP is supported by P30AG10161, P30AG72975, R01AG17917, R01 AG015819, U01 AG072572, and U01 AG046152.

## Appendix

### A Model details

#### A.1 Negative binomial parameterization

We use the negative binomial distribution that counts the number of successes *y* before *A* failures, with success probability *p*,

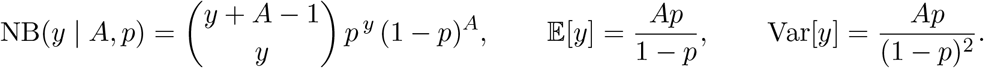

Setting *p* = *σ*(*ρ*) = 1*/*(1 + *e*^−*ρ*^) gives 1 − *p* = 1*/*(1 + *e*^*ρ*^) and *p/*(1 − *p*) = *e*^*ρ*^, so that

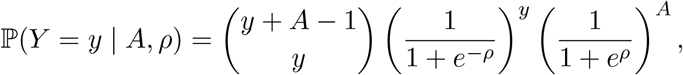

which is each term of Eq. (1), with mean *µ* = *Ae*^*ρ*^ and variance *Ap/*(1 − *p*)^2^ = *µ*(1 + *e*^*ρ*^) = *µ* + *µ*^2^*/A*. Equivalently, in the convention that counts failures before *A* successes, the success probability is 1 − *p* = (1+ *e*^*ρ*^)^−1^. Since *A* is continuous under its prior, the binomial coefficient is understood as Γ(*y* + *A*)*/* (Γ(*y* + 1)Γ(*A*)). This is the parameterization of Agrawal et al. [2026] and of NumPyro’s `NegativeBinomialLogits` with logits *ρ* and total count *A*.

#### A.2 Prior distributions

The full generative model of HiP-BIND, which combines the likelihood (1) with the priors (3) and (4), is, for *m* ∈ [*M*], *l* ∈ [*L*], *d* ∈ [*D*] and *k* ∈ {0, 1} (all draws independent unless indicated):

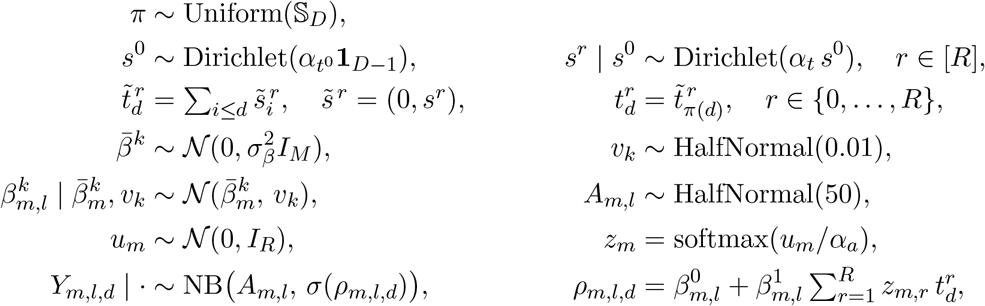

where S_*D*_ is the set of permutations of [*D*], *σ* is the logistic function, and the likelihood is evaluated only on observed entries.

The priors on *β*^0^, *β*^1^ pool information across regions (or layers) within each feature, in the spirit of the hierarchical model of Agrawal et al. [2026]: the region-specific intercepts and slopes of feature *m* are shrunk toward a feature-level mean 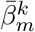 with a shared variance *v*_*k*_, which has a half-normal prior of scale 0.01. This is a simplified version of the B-BIND hierarchical prior, which pools only the slopes and infers a full covariance across layers. The feature-level means have a weakly informative prior with 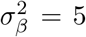 on real data and 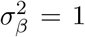 in simulations. Since 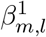 is unconstrained in sign, features may increase or decrease along the ordering. The overdispersion parameters *A*_*m,l*_ *>* 0 have a half-normal prior of scale 50. The hyperparameters are the number of groups *R*, the temperature *α*_*a*_, the concentrations 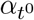 and *α*_*t*_ (values in Appendix B.1), and, for HiP-BIND-1, the Sinkhorn temperature *τ* and number of iterations 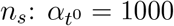 makes the master sequence close to a uniform grid, and *α*_*t*_ controls how far each cluster warp deviates from it (Fig. 5).

**Figure 5:**
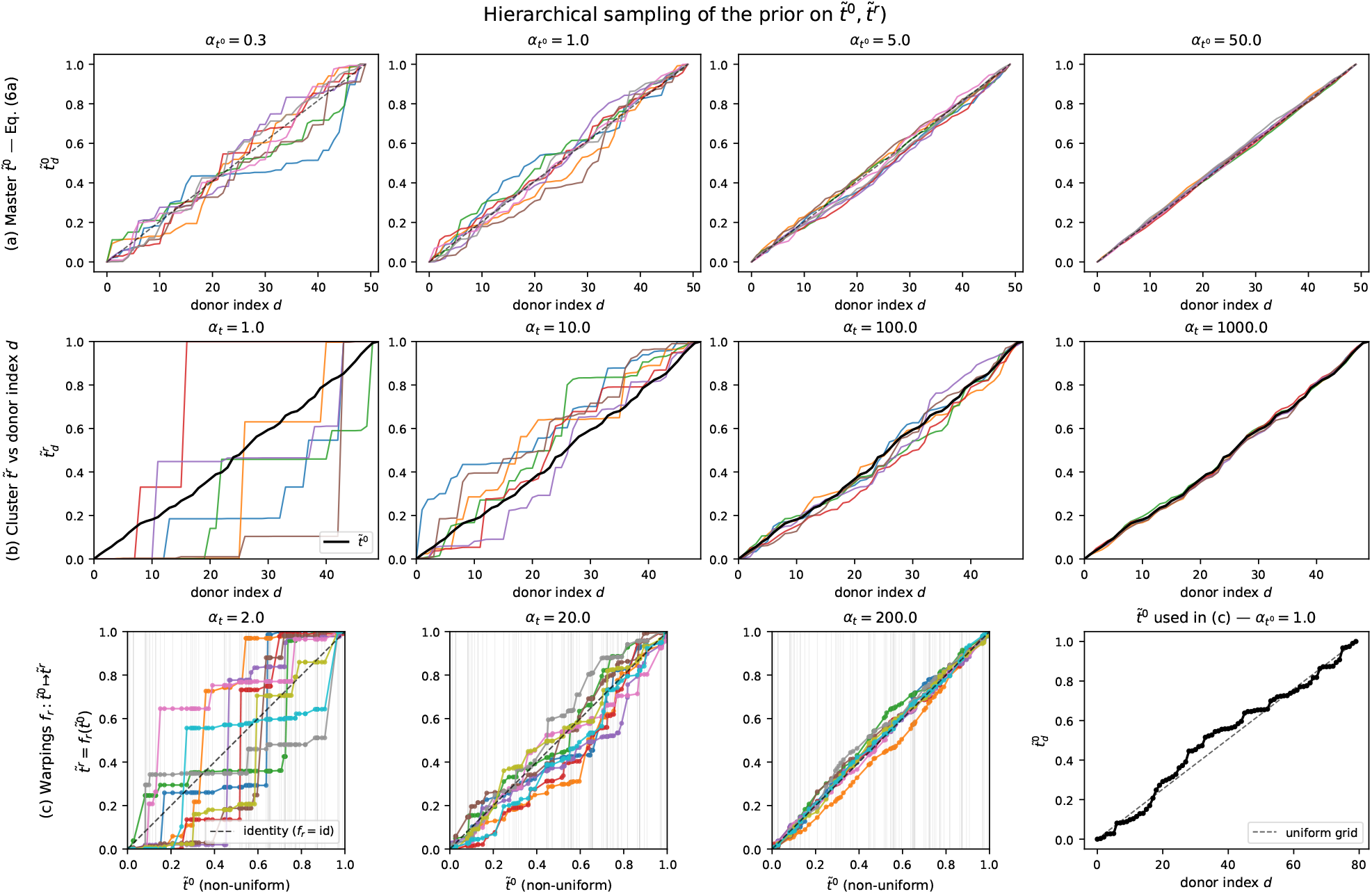
Examples of the hierarchical sampling procedure for the prior on 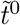 and 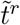 Eqs. (4a),(4b). **(a)** Master pseudotime samples 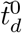 obtained from 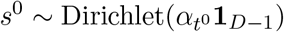 and 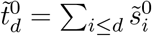, with 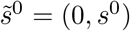, for *D* = 50 donors and four values of the dispersion 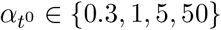 (8 draws each). Small 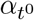 concentrates increment mass on a few coordinates and produces staircase-like sequences; large 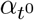 collapses 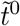 onto the uniform grid *d/*(*D* − 1) (dashed). **(b)** Cluster-specific sequences 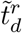 drawn conditional on a single fixed master draw 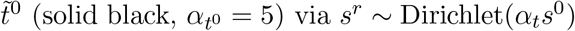, shown for four concentration levels *α*_*t*_ ∈ {1, 10, 100, 1000} (6 draws each). The conditional Dirichlet has mean *s*^0^, so 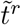 is centered on 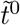 with concentration controlled by *α*_*t*_: large *α*_*t*_ pulls 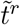 toward 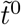, while small *α*_*t*_ yields broad excursions concentrated on a few donors. **(c)** The same construction visualized as the implicit monotone warping 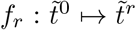, plotted against 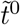 (no longer the donor index) for a deliberately non-uniform master clock (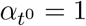, *D* = 80). The dashed diagonal marks the identity *f*_*r*_ = id; light gray verticals indicate the sampled 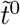 values (*x*-axis stretching reflects the non-uniformity). The right tile shows the underlying 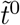 used in the three warping panels. As *α*_*t*_ grows, *f*_*r*_ concentrates on the identity; as it shrinks, *f*_*r*_ fans out into a family of monotone, endpoint-pinned warpings. Together, panels **(a)**–**(c)** illustrate how the prior decouples the shape of the master clock (controlled by 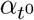) from the spread of cluster-specific deviations around it (controlled by *α*_*t*_).

For a controlled comparison, we implement B-BIND within the same framework as HiP-BIND: it uses the B-BIND likelihood and pseudotime prior, *t*_*d*_ ~ Beta(1, 1) i.i.d., with the same priors on *β*^0^, *β*^1^ and *A* and the same SVI procedure as HiP-BIND.

For a controlled comparison, we implement B-BIND with the same likelihood, coefficient and overdispersion priors, and SVI procedure as HiP-BIND, while retaining its independent pseudotime prior, *t*_*d*_ ~ Beta(1, 1). Thus, *R* = 1 HiP-BIND and B-BIND are closely matched single-trajectory models.

#### A.3 Hierarchical prior on the pseudotimes

### B Experimental setup and model selection

#### B.1 Hyperparameters and inference

For the simulations in Section 3 we used: *R* ∈ {1, 2, 3, 4, 5}, *α*_*a*_ = 0.05, 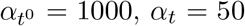, *α*_*t*_ = 50 and 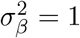, with the AutoNormal guide and the Adam optimizer with learning rate 0.0005. B-BIND was fit for 20,000 SVI steps with 10 ELBO particles. HiP-BIND-0 was initialized at the B-BIND ordering and alternated 5 blocks of 20,000 SVI steps with Hungarian updates of *P*, followed by a final 20,000-step refinement, with 20 ELBO particles. HiP-BIND-1 was initialized at the HiP-BIND-0 solution and fit for 20,000 steps with 20 ELBO particles, Sinkhorn temperature *τ* = 0.01 and *n*_*s*_ = 100 Sinkhorn iterations. For PCA, the ordering is given by the first principal component of the standardized features. Posterior summaries use 2,000 samples from the fitted guide.

In HiP-BIND-1, the Gumbel-Sinkhorn prior on *B*(*D*) is set by iteratively applying the Sinkhorn algorithm (i.e., iteratively scaling the rows and columns) to an elementary random matrix with Gumbel entries. Since this iterative scaling induces a reparameterization of *P* as a differentiable transformation of a noise distribution, we can perform variational inference. In the limit of infinite Sinkhorn iterations, this matrix is guaranteed to be doubly stochastic [Cominetti and Martín, 1994, Cuturi, 2013] although it may fail to be a *bona-fide* permutation. The temperature *τ* and the number of Sinkhorn iterations *n*_*s*_ are controllable parameters.

For the real-data experiments in Section 4 we considered the following parameter configurations: *R* ∈ {1, 2, 3, 4, 5}, *α*_*a*_ ∈ {0.01, 0.05, 0.1, 0.5}, 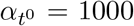 (we chose such a large value so that the master sequence 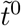 is close to a uniform grid), *α*_*t*_ ∈ {20, 100, 200}. For each of these choices we performed stochastic variational inference as implemented in NumPyro using the AutoNormal guide with the Adam optimizer, with learning rate 0.005 and 10 ELBO particles. For each configuration we ran 21 random seeds for ROSMAP and 4 for SEA-AD, where in HiP-BIND-0 the seed also sets the initial permutation *P*. For HiP-BIND-1 we used the final means given by HiP-BIND-0 as initializations.

#### B.2 Selection of runs

Different seeds and hyperparameter configurations lead to distinct solutions, indicating a multimodal variational objective. For each fitted run we compute a label-free self-consistency score: the mean over features of the absolute Spearman correlation between each layer- or region-averaged observed feature and the inferred donor ordering. The score uses neither staging labels nor, in simulations, the true ordering. For the real-data results, we rank all fitted (configuration, seed) runs by this score and report averages over the 20 highest-ranked runs. This discards degenerate solutions whose ordering is driven by a few features. Since the sign of 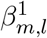 is unconstrained, reversing the ordering and flipping the signs of *β*^1^ leaves the likelihood unchanged. We therefore report absolute Spearman correlations throughout.

Applying the same selection to B-BIND has no effect: all B-BIND seeds converge to essentially the same solution (scores within 0.0005 in ROSMAP and 0.002 in SEA-AD; pairwise |Spearman| ≥ 0.999 between orderings). B-BIND has no counterpart of the HiP-BIND grid over (*R, α*_*a*_, *α*_*t*_), and since its seeds converge to the same solution, additional restarts would not change it. Conversely, the advantage of HiP-BIND does not depend on the selection: the median over all *R* = 3 runs, without any selection, attains Braak/Thal/ADNC/CERAD correlations of 0.745/0.837/0.884/0.831 in ROSMAP and 0.684/0.830/0.790/0.639 in SEA-AD, compared with 0.648/0.844/0.828/0.816 and 0.650/0.828/0.769/0.625 for B-BIND. Across all *R* = 3 runs, HiP-BIND exceeds the B-BIND average on Braak, ADNC and CERAD in 92%, 88% and 68% of runs in ROSMAP and in 86%, 83% and 91% in SEA-AD; moreover, 71% (ROSMAP) and 76% (SEA-AD) of all HiP-BIND runs attain a higher score than the best B-BIND seed.

#### B.3 Sensitivity to the number of latent trajectories

We fit HiP-BIND with *R* = 1, …, 5 and score every run with the self-consistency criterion of Appendix B.2. Because additional trajectory groups can only increase flexibility, the score typically rises sharply up to an adequate *R* and is flat beyond it. We use this pattern to detect an under-specified *R* and to check how much the results depend on *R* beyond it. In simulations, where the true *R* is known, we also check whether it identifies *R*: we select the smallest *R* whose score is within 0.01 of its maximum over *R* ∈ {1, …, 5}.

For each *R* we report the self-consistency score and the number of occupied groups. Each feature is assigned to its most probable group, arg max_*r*_ *z*_*m,r*_, and a group is occupied if at least one feature is assigned to it; since the model may leave groups empty, this number can be smaller than *R*. We write “*k* (*n*)” when *n* runs (real data) or datasets (simulations) use *k* occupied groups. In simulations, where the truth is known, we also report the Spearman correlation between the inferred and the true ordering, the adjusted Rand index (ARI) between the inferred and the true feature groups, and the number of datasets in which the selection rule chooses each *R*.

##### Simulations

For each of the 20 simulated datasets of Section 3, we fit HiP-BIND-0 with *R* = 1, …, 5 using the settings of Appendix B.1, initializing the ordering from the B-BIND solution. Table 3 reports the Spearman correlation with the true ordering, the self-consistency criterion, the adjusted Rand index (ARI) of the inferred feature groups, the number of occupied groups, and how often each *R* is selected by the rule above. Under-specifying *R* is harmful: moving from *R* = 1 to *R* = 2 increases the Spearman correlation by 0.238 (paired 95% CI [0.220, 0.256]; better in all 20 datasets). Over-specifying *R* leaves the recovered ordering essentially unchanged (differences from *R* = 2 are at most 0.003), but the surplus groups split the true feature groups, which lowers the ARI. The criterion rises sharply from *R* = 1 to *R* = 2 and is flat thereafter, so its plain maximum often falls at *R* = 3 or *R* = 4 (17 of 20 datasets), whereas the selection rule, which favors the smallest *R* within 0.01 of the maximum, recovers the true *R* = 2 in all 20 datasets. The *R* = 2 fits were rerun for this analysis and differ from HiP-BIND-0 in Table 1 only through numerical variation between runs (mean Spearman 0.969 vs. 0.967).

**Table 3:** Simulation results for *R* = 1, …, 5 over 20 datasets (true *R* = 2): mean ± s.d. of the Spearman correlation with the true ordering, of the self-consistency score and of the ARI; occupied groups; and number of datasets in which the selection rule chooses each *R*.

| Model | Spearman | Self-consistency | ARI | Occupied groups | Selected |
| --- | --- | --- | --- | --- | --- |
| B-BIND | $0.729 \pm 0.045$ | – | – | 1 (20) | – |
| $R = 1$ | $0.731 \pm 0.040$ | $0.587 \pm 0.017$ | – | 1 (20) | 0/20 |
| $R = 2$ | $0.969 \pm 0.008$ | $0.669 \pm 0.012$ | $0.88 \pm 0.19$ | 2 (20) | 20/20 |
| $R = 3$ | $0.966 \pm 0.007$ | $0.671 \pm 0.013$ | $0.46 \pm 0.09$ | 3 (20) | 0/20 |
| $R = 4$ | $0.968 \pm 0.009$ | $0.670 \pm 0.012$ | $0.66 \pm 0.15$ | 3 (9), 4 (11) | 0/20 |
| $R = 5$ | $0.971 \pm 0.009$ | $0.669 \pm 0.012$ | $0.82 \pm 0.00$ | 3 (20) | 0/20 |

**Table 4:** Real-data results for *R* = 1, …, 5, averaged over the 20 highest-ranked runs at each *R* (the *R* = 3 rows coincide with Table 2): self-consistency score, |Spearman| with the staging scores (all donors), and number of occupied groups across these runs.

| Cohort | Model | Self-consistency | Braak | Thal | ADNC | CERAD | Occupied groups (runs) |
| --- | --- | --- | --- | --- | --- | --- | --- |
| ROSMAP | B-BIND | 0.765 | 0.648 | 0.844 | 0.828 | 0.816 | 1 |
| | $R = 1$ | 0.766 | 0.648 | 0.845 | 0.829 | 0.820 | 1 (20) |
| | $R = 2$ | 0.784 | 0.745 | 0.842 | 0.885 | 0.836 | 2 (20) |
| | $R = 3$ | 0.784 | 0.740 | 0.842 | 0.883 | 0.836 | 2 (18), 3 (2) |
| | $R = 4$ | 0.786 | 0.728 | 0.848 | 0.879 | 0.834 | 2 (17), 3 (3) |
| | $R = 5$ | 0.784 | 0.748 | 0.840 | 0.886 | 0.834 | 2 (18), 3 (2) |
| SEA-AD | B-BIND | 0.750 | 0.650 | 0.828 | 0.769 | 0.625 | 1 |
| | $R = 1$ | 0.752 | 0.655 | 0.829 | 0.772 | 0.630 | 1 (20) |
| | $R = 2$ | 0.762 | 0.726 | 0.828 | 0.822 | 0.688 | 2 (20) |
| | $R = 3$ | 0.760 | 0.710 | 0.843 | 0.814 | 0.665 | 2 (11), 3 (9) |
| | $R = 4$ | 0.760 | 0.715 | 0.839 | 0.817 | 0.665 | 2 (10), 3 (10) |
| | $R = 5$ | 0.762 | 0.732 | 0.846 | 0.828 | 0.677 | 2 (12), 3 (8) |

##### Real data

For each cohort and each *R* ∈ {1, …, 5} we rank all fitted configurations and seeds by the self-consistency score and, to obtain a more stable summary across *R*, average the 20 highest-ranked runs at each *R*. The pattern observed in the simulations recurs: *R* = 1 attains the same score as B-BIND, the score increases at *R* = 2, and it is flat thereafter. Because the score differences are about an order of magnitude smaller than in the simulations (+0.018 in ROSMAP and +0.010 in SEA-AD from *R* = 1 to *R* = 2), a fixed tolerance calibrated in simulations does not transfer, and we compare runs directly instead. Every one of the 20 highest-ranked runs with *R ≥* 2 has a higher self-consistency score than the best run with *R* = 1 in both cohorts, whereas the mean scores for *R* = 2, …, 5 differ by at most 0.002. The staging correlations follow the same pattern: *R* = 1 matches B-BIND, and *R* = 2, …, 5 give similar values, all above B-BIND on Braak, ADNC and CERAD. The fitted models use only two or three occupied groups for every *R ≥* 2, leaving additional groups empty even for *R* = 4, 5. The choice among *R ≥* 2 therefore matters little. We report *R* = 3 in the main text, which accommodates the up to three groups used by the fits. This is also motivated by biological reasons: it allows for amyloid- and tau-dominant groups plus a third group for mixed or region-specific dynamics, such as the medial temporal tau pattern in Fig. 12, without forcing one.

### C Computational complexity and runtime

A single-pseudotime model (B-BIND) has *O*(*D*) time parameters, whereas HiP-BIND represents the ordering by a *D* × *D* matrix *P*, requiring *O*(*D*^2^) memory; the *R* cluster sequences add *O*(*RD*) parameters. Each SVI step evaluates the likelihood at cost *O*(*DML*) for both models and, for HiP-BIND, applies *P* to the *R* + 1 increasing sequences at cost *O*(*RD*^2^). In HiP-BIND-0, the permutation update is performed about ten times per fit, outside the SVI loop: building the *D* × *D* cost matrix of Section 2.2 costs *O*(*D*^2^*ML*) and the Hungarian solve costs *O*(*D*^3^); one full update takes 3.2 s at *D* = 1218 (0.04 s at *D* = 84), negligible relative to SVI. HiP-BIND-1 instead unrolls *n*_*s*_ Sinkhorn iterations inside every gradient step, at *O*(*n*_*s*_*D*^2^) time and memory per step.

Table 5 compares the models under the inference settings used in the paper (AutoNormal guide, Adam with learning rate 0.005, 10 ELBO particles). Per SVI step, HiP-BIND-0 costs 1.3× (ROSMAP) and 1.4× (SEA-AD) the B-BIND step, and this cost is nearly independent of the number of clusters (19.1 vs. 21.2 ms on ROSMAP for *R* = 1 and *R* = 3). A full HiP-BIND-0 fit takes 5.6× (ROSMAP: 31.4 vs. 5.6 min) and 6.8× (SEA-AD: 177 vs. 26 s) longer than a B-BIND fit, mainly because it runs more SVI iterations (72.5k vs. 20k, including warm restarts between permutation updates), not because each step is more expensive. The additional cost is therefore due mostly to the alternating optimization of the shared ordering rather than to the cluster structure itself; in absolute terms, a full fit takes at most about half an hour on a laptop CPU, and fits are independent across seeds and configurations, so they parallelize trivially. HiP-BIND-1 scales quadratically in *D* per step: a single 20k-step fit on ROSMAP would take about 14 h. For this reason, all real-data results use HiP-BIND-0.

**Table 5:** Computational cost on an 8-core Apple M2 CPU (full cohorts; AutoNormal guide, Adam with learning rate 0.005, 10 ELBO particles). Per-step times: steady-state SVI throughput (JIT compilation excluded). Full-fit times: end-to-end wall-clock for one fit (B-BIND: 20k SVI steps; HiP-BIND-0: 72.5k steps with 10 permutation updates; HiP-BIND-1: 20k steps, *n*_*s*_ = 5 Sinkhorn iterations). ^*†*^Extrapolated from the per-step time; all other full-fit times are measured. –: not applicable.

| Cohort | Measure | B-BIND | HiP-BIND-0 | HiP-BIND-1 |
| --- | --- | --- | --- | --- |
| SEA-AD ($D = 84$) | per SVI step | 0.95 ms | 1.34 ms | 5.85 ms |
|  | permutation update | – | 0.04 s | – |
|  | full fit | 26 s | 177 s | 138 s |
| ROSMAP ($D = 1218$) | per SVI step | 16.2 ms | 21.2 ms | 2.5 s |
|  | permutation update | – | 3.2 s | – |
|  | full fit | 5.6 min | 31.4 min | 14 h<sup>†</sup> |

These timings are for a single fit on a laptop CPU. The experiments in the paper were run as independent single-fit jobs on parallel CPUs of a high-performance cluster. The accompanying code allows one to replicate a few representative seeds for the SEA-AD dataset, as well as the simulated scenarios. For this additional cost, HiP-BIND-0 yields statistically significant gains over B-BIND on Braak, ADNC and CERAD in both cohorts (Appendix D.2). The real-data results select among many fits, which at the timings of Table 5 amounts to about 130 CPU-hours for ROSMAP. This search is not needed for the gain itself: a single randomly chosen *R* = 3 fit, at 5.6–6.8× the cost of a B-BIND fit, already exceeds B-BIND on Braak and ADNC in 83–92% of cases (Appendix B.2); the selection mainly removes degenerate solutions and reduces run-to-run variability.

### D Statistical significance and stability

#### D.1 Simulations

Each of the 20 simulation replicates is an independent dataset drawn from the scenario of Section 3, on which every method is refit and scored against the true ordering. Paired by dataset, the differences in Spearman correlation therefore reflect both data and fitting variability. Table 6 reports the mean paired difference with a paired-*t* 95% confidence interval. Both HiP-BIND variants improve on B-BIND and on PCA in every replicate. HiP-BIND-1 is initialized at the HiP-BIND-0 solution, which explains why the two perform almost identically.

#### D.2 Real data

Table 7 reports the standard deviations of the correlations in Table 2 across runs.

**Table 6:** Paired differences in Spearman correlation with the true ordering over 20 runs.

| Contrast | Mean difference | 95% CI | Datasets improved |
| --- | --- | --- | --- |
| HiP-BIND-0 – B-BIND | +0.238 | [0.216, 0.259] | 20/20 |
| HiP-BIND-0 – PCA | +0.072 | [0.064, 0.081] | 20/20 |
| HiP-BIND-1 – B-BIND | +0.238 | [0.216, 0.260] | 20/20 |
| HiP-BIND-1 – HiP-BIND-0 | 0.000 | [−0.002, 0.002] | 13/20 |

**Table 7:**
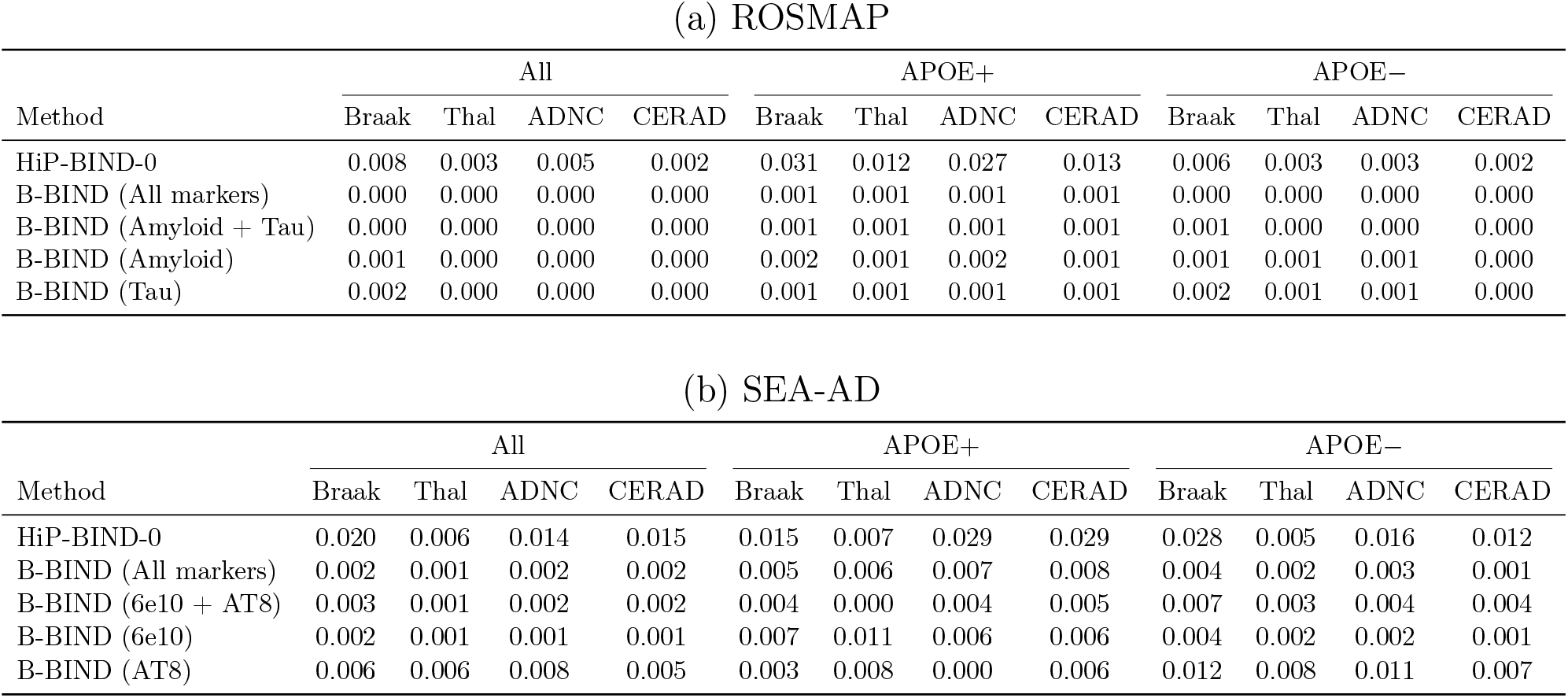
Standard deviation of the Spearman correlation between model-inferred donor orderings and standard ordinal clinical scores (Braak, Thal, ADNC, and CERAD), stratified by APOE carrier status (All, APOE+, APOE−). Standard deviations are computed across 20 independent runs.

For each cohort, we draw 2,000 bootstrap samples of donors with replacement. On each sample, we recompute |*ρ*| between each fitted ordering and the staging score, average over the selected HiP-BIND-0 runs and over the B-BIND seeds, and take the difference Δ between HiP-BIND-0 and B-BIND (All markers). The fitted orderings are held fixed, since refitting on a bootstrap sample is ill-defined when duplicated donors must occupy distinct positions of a permutation; the intervals therefore quantify donor sampling variability given the fits. Resampling the runs in addition to the donors changes the interval endpoints by at most 0.01 and does not change which intervals exclude 0. Table 8 shows that the gains on Braak, ADNC and CERAD are significant in both cohorts, whereas the differences on Thal are not. The same holds within APOE strata in ROSMAP (e.g., Braak Δ = +0.184, 95% CI [0.136, 0.242], among the 279 carriers); in SEA-AD, the strata are small (25 carriers and 59 non-carriers), and only Braak and ADNC among non-carriers reach significance. Moreover, every selected HiP-BIND-0 run has a higher correlation than every B-BIND seed for Braak, ADNC and CERAD in both cohorts.

**Table 8:** Mean |Spearman| with staging scores for B-BIND (All markers) and HiP-BIND-0 in the full cohorts, their difference Δ (computed before rounding), the 95% donor-bootstrap percentile interval of Δ (2,000 resamples, fitted orderings held fixed), and the percentage of bootstrap resamples in which HiP-BIND-0 is higher.

| Cohort | Score | B-BIND | HiP-BIND-0 | $\Delta$ | 95% CI | Resamples improved |
| --- | --- | --- | --- | --- | --- | --- |
| SEA-AD<br>($D = 84$) | Braak | 0.650 | 0.710 | +0.060 | [0.022, 0.109] | 100% |
|  | Thal | 0.828 | 0.843 | +0.015 | [−0.003, 0.040] | 94.6% |
|  | ADNC | 0.769 | 0.814 | +0.046 | [0.014, 0.090] | 99.9% |
|  | CERAD | 0.625 | 0.665 | +0.041 | [0.006, 0.085] | 99.2% |
| ROSMAP<br>($D = 1218$) | Braak | 0.648 | 0.740 | +0.092 | [0.074, 0.110] | 100% |
|  | Thal | 0.844 | 0.842 | −0.002 | [−0.013, 0.009] | 35.8% |
|  | ADNC | 0.828 | 0.883 | +0.055 | [0.044, 0.068] | 100% |
|  | CERAD | 0.816 | 0.836 | +0.020 | [0.009, 0.032] | 100% |

The orderings of the selected HiP-BIND-0 runs agree closely: the mean pairwise |Spearman| between runs is 0.989 in ROSMAP (minimum 0.977) and 0.990 in SEA-AD (minimum 0.984), and the median over donors of the standard deviation of a donor’s rank across runs is 1.6% (ROSMAP) and 1.3% (SEA-AD) of *D*, smallest for early-stage donors. B-BIND converges to essentially the same ordering from every seed (pairwise |Spearman| ≥ 0.999).

These quantities describe variability across runs and donor resamples, not calibrated posterior intervals. In HiP-BIND-0 we use a small Gumbel noise scale, so each permutation update acts as a MAP step for *π* with random tie-breaking, and each fit returns a point estimate of the ordering. HiP-BIND-1 provides a relaxed posterior over *P* (Fig. 6), but variational approximations tend to underestimate posterior uncertainty; assessing the calibration of the resulting intervals is left for future work.

**Figure 6:**
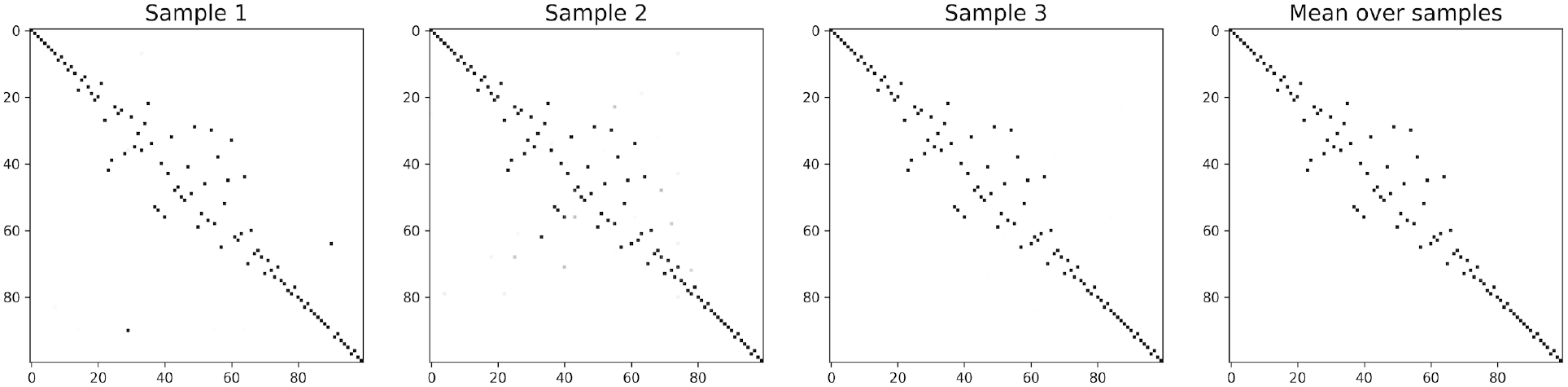
Samples of *P* from the inferred HiP-BIND-1 for the simulation setup described in 3. The last column shows the mean across samples. The samples differ in only a few entries, indicating a concentrated approximate posterior.

### E Additional simulations results

Fig. 6 shows samples of the relaxed permutation matrix *P* from the HiP-BIND-1 variational posterior in the scenario of Section 3, with donors in their true order, so that a diagonal matrix corresponds to exact recovery. Early and late donors are ordered almost exactly, whereas deviations from the diagonal concentrate in the middle of the progression. The samples differ in only a few entries, so the approximate posterior is concentrated around a single ordering; as noted in Appendix D.2, this spread should not be read as calibrated uncertainty.

Fig. 7 and Table 9 consider three further pairs of warping functions for the two feature groups. These scenarios are less challenging for a single pseudotime than that of Section 3: all methods recover the donor ordering well (Spearman 0.91–0.95), the differences between methods are at most 0.018, and HiP-BIND-0 is best in the exponential versus power-law scenario and PCA in the other two. The difference lies in the trajectories rather than in the ordering: HiP-BIND-0 recovers both warps in all three scenarios, whereas the PCA trajectories are not monotone and B-BIND underestimates the late rise of the power-law group in the first scenario.

**Table 9:** Spearman correlation and Mallows distance for the three simulation scenarios of Fig. 7, averaged across 20 independent runs. The best mean per scenario is in bold.

| Experimental Setup | Method | Spearman (mean ± std) | Mallows (mean ± std) |
| --- | --- | --- | --- |
| exp vs power | PCA | 0.9163 ± 0.0166 | 11.7511 ± 1.2472 |
| | B-BIND | 0.9271 ± 0.0220 | 10.9047 ± 1.6465 |
| | HiP-BIND-0 | **0.9309** ± 0.0206 | **10.6239** ± 1.5643 |
| | HiP-BIND-1 | 0.9286 ± 0.0230 | 10.7757 ± 1.7363 |
| linear vs power | PCA | **0.9140** ± 0.0245 | **11.8528** ± 1.7006 |
| | B-BIND | 0.9093 ± 0.0263 | 12.1735 ± 1.7636 |
| | HiP-BIND-0 | 0.9068 ± 0.0261 | 12.3472 ± 1.7115 |
| | HiP-BIND-1 | 0.9137 ± 0.0251 | 11.8678 ± 1.7478 |
| quadratic vs hill | PCA | **0.9473** ± 0.0118 | **9.3193** ± 1.0367 |
| | B-BIND | 0.9294 ± 0.0126 | 10.8038 ± 0.9604 |
| | HiP-BIND-0 | 0.9390 ± 0.0140 | 10.0237 ± 1.1383 |
| | HiP-BIND-1 | 0.9390 ± 0.0139 | 10.0187 ± 1.1360 |

**Figure 7:**
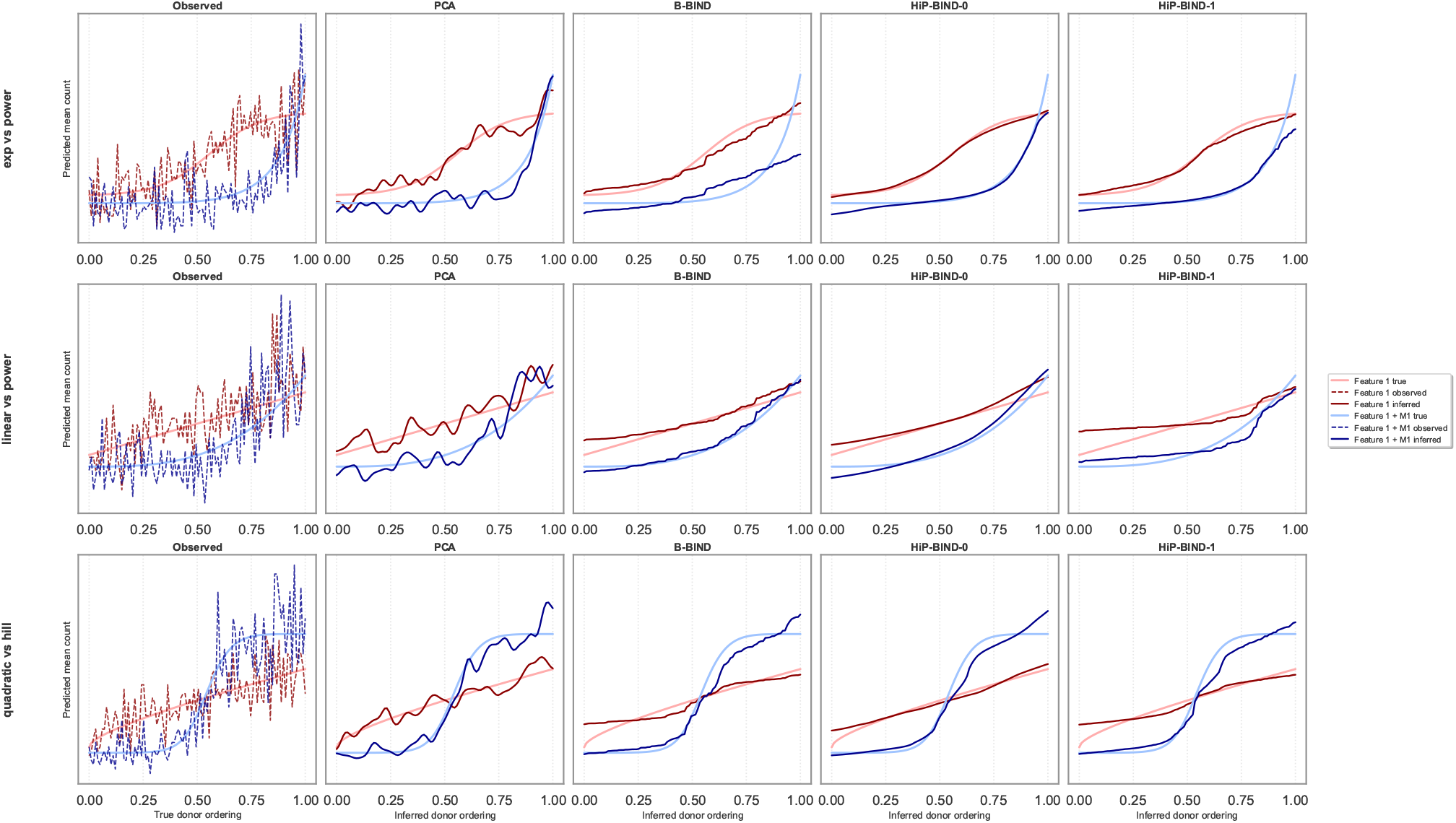
Simulation scenarios illustrating recovery of heterogeneous temporal dynamics under different latent warping functions. Rows correspond to three synthetic settings with distinct pairs of temporal processes: exponential versus power-law (top), linear versus power-law (middle), and quadratic versus Hill-type dynamics (bottom). Columns compare the observed data and trajectories inferred by PCA, B-BIND, HiP-BIND-0, and HiP-BIND-1. Red and blue curves denote the two latent feature groups.

To assess robustness to features that carry no information about the ordering, we simulate 2 signal features, whose means follow the same sigmoid-shaped rise along the true ordering, among 10, 20 or 40 features in total; the remaining features have constant means (Fig. 8). With 10 features, B-BIND and both HiP-BIND variants perform similarly (Spearman 0.85–0.86) and PCA slightly worse (0.82; Table 10). As the number of noise features grows, the Spearman correlation of B-BIND drops from 0.85 to 0.34 and that of PCA from 0.82 to 0.56, whereas both HiP-BIND variants stay above 0.80. The fitted trajectories show the same pattern (Fig. 9): with 40 features, the B-BIND signal trajectory rises only gradually and misses the sigmoid shape, and the PCA trajectory is not monotone, whereas HiP-BIND still recovers the rise of the signal features, although neither variant reproduces its final plateau exactly. B-BIND and HiP-BIND fit the noise features as flat. With 40 features HiP-BIND-0 varies more across runs (s.d. 0.14) than HiP-BIND-1 (0.05), which is initialized at the HiP-BIND-0 solution.

**Table 10:** Spearman correlation and Mallows distance with 2 signal features among 10, 20 and 40 features in total (the rest are pure noise), averaged across 20 independent runs. The best mean per setting is in bold.

| Experimental Setup | Method | Spearman (mean ± std) | Mallows (mean ± std) |
| --- | --- | --- | --- |
| 2 / 10 | PCA | 0.8156 ± 0.0205 | 17.5051 ± 0.9780 |
| | B-BIND | 0.8521 ± 0.0200 | 15.6625 ± 1.0672 |
| | HiP-BIND-0 | **0.8559** ± 0.0223 | **15.4526** ± 1.2091 |
| | HiP-BIND-1 | 0.8558 ± 0.0221 | 15.4586 ± 1.2086 |
| 2 / 20 | PCA | 0.7140 ± 0.1755 | 21.2388 ± 5.1766 |
| | B-BIND | 0.6843 ± 0.1613 | 22.2846 ± 5.5672 |
| | HiP-BIND-0 | **0.8576** ± 0.0197 | **15.3686** ± 1.0740 |
| | HiP-BIND-1 | 0.8547 ± 0.0187 | 15.5274 ± 1.0071 |
| 2 / 40 | PCA | 0.5579 ± 0.2035 | 26.5268 ± 5.8966 |
| | B-BIND | 0.3373 ± 0.1774 | 32.8894 ± 4.8836 |
| | HiP-BIND-0 | 0.8050 ± 0.1446 | 17.3035 ± 5.1799 |
| | HiP-BIND-1 | **0.8289** ± 0.0542 | **16.7200** ± 2.4280 |

**Figure 8:**
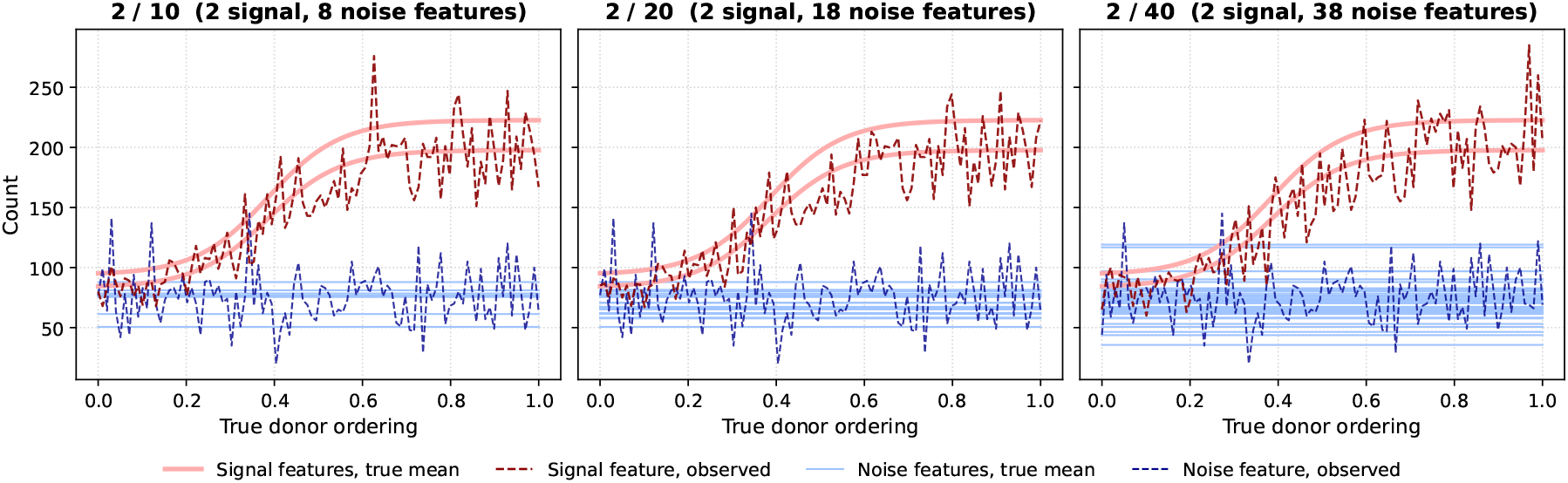
Simulated data with 2 signal features among 10, 20 and 40 features in total. Red: signal features, whose mean rises along the true donor ordering; blue: noise features with a constant mean. Solid lines are true means, dashed lines observed counts of one feature of each kind.

**Figure 9:**
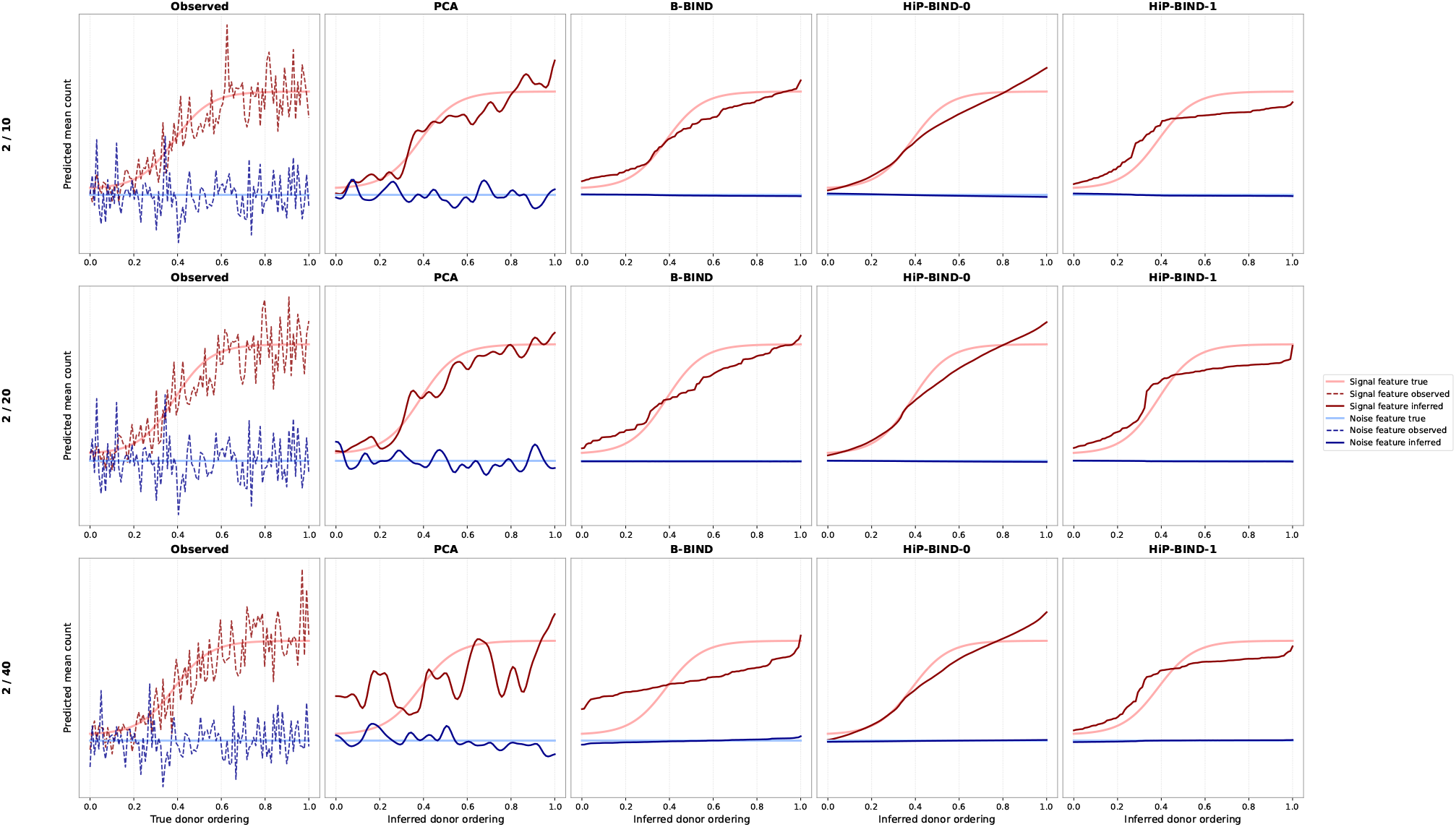
Recovery of the signal under increasing numbers of noise features. Rows correspond to 2 signal features among 10 (top), 20 (middle) and 40 (bottom) features. Columns compare the observed data and trajectories inferred by PCA, B-BIND, HiP-BIND-0, and HiP-BIND-1. Red and blue curves denote a signal and a noise feature.

### F Data access, preprocessing and features

#### Data access and inclusion

SEA-AD data are publicly available from the Allen Institute for Brain Science under its terms of use. ROSMAP data are available through the Rush Alzheimer’s Disease Center Research Resource Sharing Hub (https://www.radc.rush.edu/). Access is controlled and requires an approved request and a Data Use Agreement. For ROSMAP, we include the 1218 donors with complete pathology and staging data, a postmortem interval of at most 24 h, and no diagnosis of other dementia or Lewy body disease (from 4003 participants in the data release and 1645 with complete data).

#### Mapping to counts

The negative binomial likelihood requires integer-valued observations, so continuous measurements are mapped to a count scale before fitting, as in Agrawal et al. [2026]. In ROSMAP, all measurements are multiplied by 5 and truncated to integers, so that one unit corresponds to 0.2 on the square-root scale; in SEA-AD, each feature is rescaled so that its maximum across donors equals 50 and then rounded. In both cohorts this resolution is small relative to the variability between donors (Figs. 3, 10), so the discretization has little effect on the fitted orderings.

**Figure 10:**
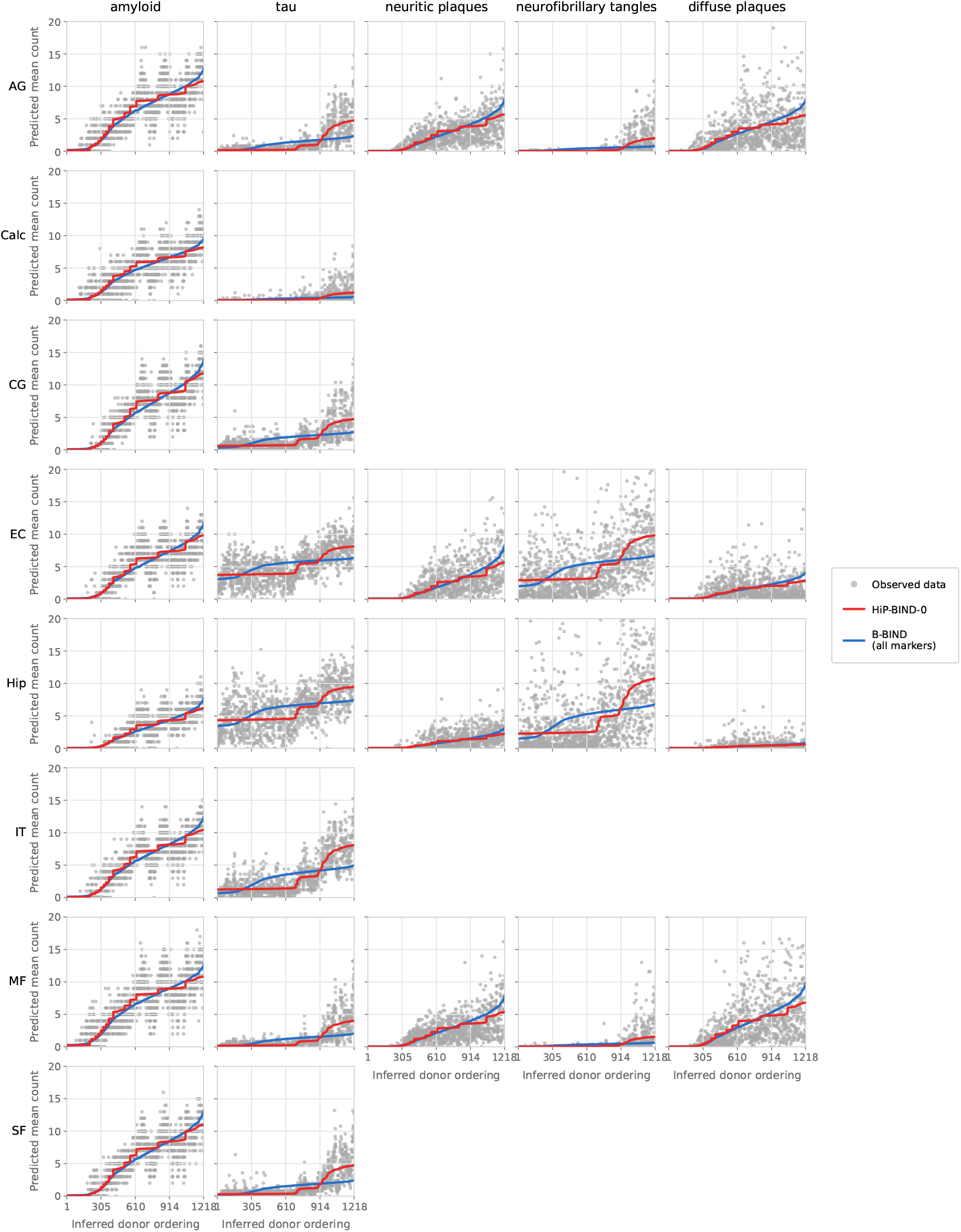
Feature-specific trajectory reconstruction across brain regions in ROSMAP. Predicted monotonic feature trajectories plotted against each model’s inferred donor ordering across all brain regions, with observed donor measurements overlaid using the HiP-BIND-0 inferred ordering. The regions are: Angular gyrus (AG), Calcarine cortex (Calc), Cingulate gyrus (CG), Entorhinal cortex (EC), Hippocampus (Hip), Inferior temporal cortex (IT), Midfrontal cortex (MF), Superior frontal cortex (SF). Gray dots represent raw cross-sectional donor measurements; red lines denote HiP-BIND-0 posterior mean trajectories, and blue lines denote the all-marker B-BIND baseline. For each dataset–model combination, the plotted run is the seed whose Spearman correlation is closest to the median across 20 seeds. Empty panels correspond to feature–region combinations not observed in the dataset. HiP-BIND captures heterogeneous progression patterns across both features and brain regions.

#### Features

Tables 11 and 12 list the features used for SEA-AD and ROSMAP, with the corresponding biological quantity and marker type.

**Table 11:** SEA-AD feature names, corresponding biological quantity, and marker type.

| Feature name | Biological measurement | Type of marker |
| --- | --- | --- |
| Percent 6e10 positive area | Amyloid-beta immunoreactive area | Pathological peptide |
| Number of 6e10 positive objects per area | Number of amyloid-beta plaques | Pathological peptide |
| Percent of Iba1 and 6e10 positive co-localized objects | Iba1 immunoreactive area (microglia) | Cellular-pathological colocalization |
| Number of Iba1 and 6e10 positive co-localized objects per area | Iba1 immunoreactive area (microglia) | Cellular-pathological colocalization |
| Average 6e10 positive object area | Size of amyloid-beta plaques | Pathological peptide |
| Number of AT8 positive cells per area | Number of pTau positive cells | Pathological peptide |
| Percent AT8 positive area | pTau immunoreactive area | Pathological peptide |
| Percent NeuN positive area | NeuN immunoreactive area (mature neurons) | Cellular |

**Table 12:** ROSMAP feature names, corresponding biological quantity, and marker type.

| Feature name | Biological measurement | Type of marker |
| --- | --- | --- |
| amylsqrt_est_(region) | Amyloid density after square root transformation in (region) | Pathological peptide |
| tangsqrt_est_(region) | Neurofibrillary tau tangle density after square root transformation in (region) | Pathological peptide |
| nft_(region) | Semi-quantitative summary measure of neurofibrillary tangles | Pathological peptide |
| plaq_n_(region) | Neuritic plaque burden summary in (region) | Pathological peptide |
| plaq_d_(region) | Diffuse plaque burden summary in (region) | Pathological peptide |

### G Additional Real Data results

This appendix shows the trajectories of all features and the clustering results summarized in Section 4.

#### Trajectories (Figs. 10, 11)

Both models give similar trajectories for amyloid- and plaque-related features. They differ mainly for tau and tangles: HiP-BIND-0 captures their late, sharp rise, most visible in the neocortical regions of ROSMAP, whereas B-BIND fits a more gradual increase. In SEA-AD, percent NeuN area decreases along the ordering, which the unconstrained sign of 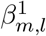 allows.

**Figure 11:**
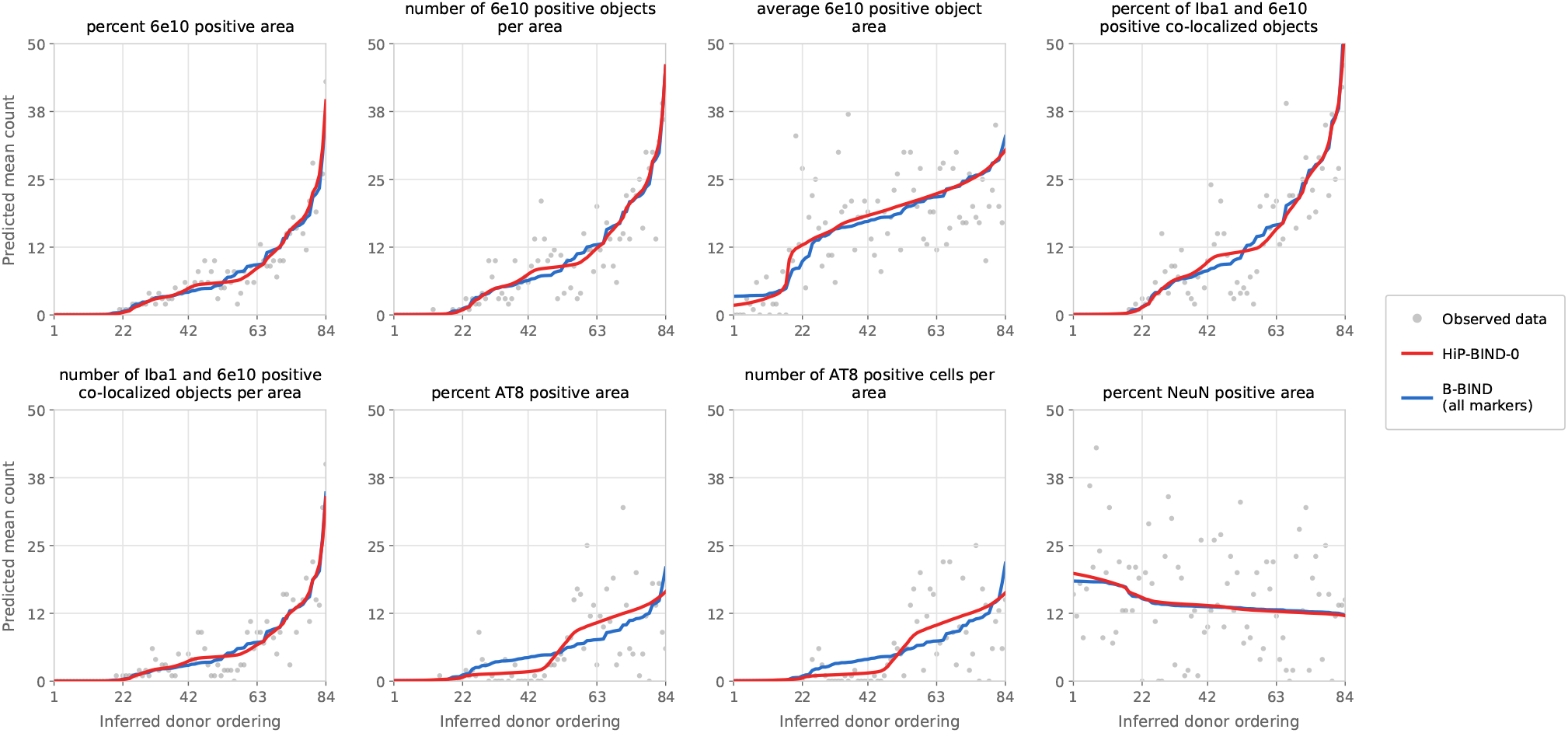
Feature-specific trajectory reconstruction in SEA-AD. Predicted monotonic feature trajectories plotted against each model’s inferred donor ordering, with observed donor measurements overlaid using the HiP-BIND-0 inferred ordering. Gray dots represent raw cross-sectional donor measurements; red lines denote HiP-BIND-0 posterior mean trajectories, and blue lines denote the all-marker B-BIND baseline. For each dataset–model combination, the plotted run is the seed whose Spearman correlation is closest to the median across 20 seeds. HiP-BIND captures heterogeneous progression patterns across features.

#### Region-resolved clustering (Fig. 12)

To test whether the same feature follows different dynamics in different regions, we refit ROSMAP with each feature-region pair as a separate feature (*M*′ = *M* × *L, L*′ = 1), restricted to EC, Hip, AG and IT, so that each pair has its own assignment. All amyloid and plaque measurements still form one cluster, whereas tau splits by region: tau and tangles from EC and Hip form a tight group, joined by IT tau at a larger distance, while AG tau and tangles form a separate group.

**Figure 12:**
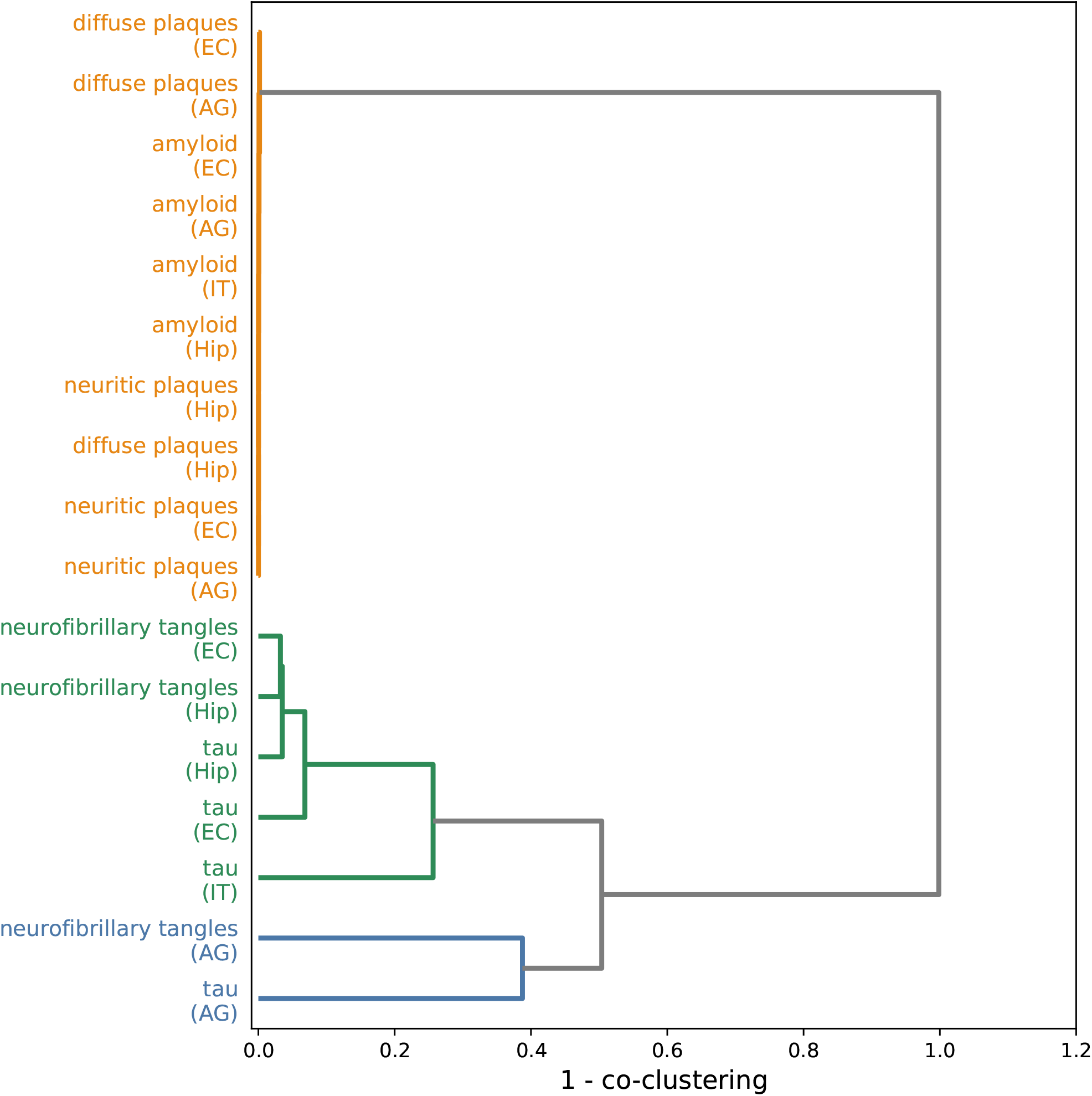
Region-resolved feature clustering for the flattened ROSMAP dataset. Dendrogram illustrating hierarchical clustering of pathology measurements when each feature–region pair is treated as a separate pseudo-feature. Distance is measured as 1 − *P* (co-cluster), where *P* (co-cluster) denotes the average soft co-clustering probability across 20 runs of HiP-BIND-0. Consistent with the higher-level clustering structure shown in Fig. 4, amyloid-related measurements across four ROSMAP brain regions (entorhinal cortex, hippocampus, angular gyrus, and inferior temporal cortex) remain grouped within a shared cluster, while tau-related measurements separate into distinct region-specific groups. In particular, tau features from the entorhinal cortex and hippocampus cluster together, reflecting the known early involvement of medial temporal lobe regions in tau pathology progression.

